# Wearable Disposable Electrotherapy

**DOI:** 10.1101/2023.11.28.569062

**Authors:** Mohamad FallahRad, Zeeshan Chaudhry, Mojtaba Belali Koochesfahani, Rayyan Bhuiyan, Mahdi Zaman, Tiffany Liu, Kisholoy Saha, Miguel R Diaz Uraga, Osvaldo Velarde, Kyle Donnery, Benjamin Babaev, Matthew Saw, Ayman Rddad, Myesha Thahsin, Alexander Couzis, Marom Bikson

## Abstract

We design and validate a novel electrotherapy platform without electronic components, using printed abundant, environmentally benign materials. Whereas existing electrotherapy devices use an independent power source and electronics to generate and control stimulation currents, our design eliminates the need for these components. Device production relies only on scalable additive manufacturing and common materials, minimizing cost and environmental impact. The disposable single-use platform (as discreet as adhesive bandages) is activated simply by placement on the body. A prescribed electrotherapy discharge is regulated by a flexible 3D electrochemical architecture tailored to each application by a novel operational theory. The single-dose usability of this platform is a categorical shift from existing approaches with durable equipment that require programming and assembly to disposable electrodes for each use. Our Wearable Disposable Electrotherapy technology can be distributed like pharmacotherapy, with indications spanning neuromodulation of brain disorders, wound healing, transcutaneous drug delivery, bioelectronic medicine, and aesthetics.

## Main

Applications of non-invasive electrical stimulation span treatment of pain and headache (1, 2), depression, addiction, age related cognitive decline (3, 4), wound healing (5, 6), aesthetic uses (7), bioelectronic medicine, and drug delivery (8). Electrical therapy devices have become compoundingly complex (microelectronics, stretchable electronics, wireless connectivity, etc.) (6, 9, 10). Battery-powered electronic devices must be connected to electrodes before each use, attached to the body, a discharge program initiated, and charged for subsequent use. The form factor of conventional devices is thus a barrier to electrotherapy healthcare adoption and compliance. The up-front-cost and cumbersomeness of electrical therapy contrasts with the “take and forget’’ usability of tablet pharmaceuticals - which contributes to the significant differences in pharmaceutical-vs electrotherapy-based healthcare (11).

In this paper, we report on the first electrotherapy platform that provides auto-initiated (upon application to the skin) controlled discharge in a single-use, disposable, low-cost, conformable bandage. Enabling these features is an integrated printable design, absent any electronics (no circuit components), with power source, self-limiting mechanism, and interface elements made from environmentally benign common materials assembled layer by layer. The entire platform is thus printed on a common substrate, with an emergent 3D electrochemical architecture that is novel and enabling. In human trials, we establish our printed battery technology, with self-limited electrical output to prescribed doses, with “apply and forget” usability. The dematerialized product is economically- and environmentally-efficient to electronics-based equipment.

The Wearable Disposable Electrotherapy platform supports scalability and distribution models akin to pharmaceuticals. Each therapy-strip can be discreetly carried, applied in any environment, and disposed after use, similar to an adhesive bandage. Indeed one application is accelerated wound healing. Adoption is seamless as caregivers (e.g. nurses, parents) are provided a product used identically to regular bandages. The potential addition of drugs with charged carriers to Wearable Disposable Electrotherapy allows a new class of transdermal drug-eluting therapy. Here, compliance compared to tablet drugs may be enhanced by continuous transdermal infusion (12). Almost any existing application of non-invasive electrotherapy can be enhanced, including democratized access to neuromodulation for pain (migraine) and neuropsychiatric disorders (3; 13).

We show the creation of Wearable Disposable Electrotherapy involved development of system design processes, a theoretical framework for self-limited dose control, novel battery cell technology, and battery pack architecture adapted to scalable manufacturing processes. For an exemplary device, in exhaustive detail, the performance of each design element is verified, and efficacy is validated in human trials. We additionally demonstrate the platform’s robustness for three applications: neuromodulation, accelerated wound healing, and iontophoresis. The impact of usability, economics, environmental, and healthcare equity are explained.

## Results and Discussion

### System design

Our goal is to develop a single-use disposable electrotherapy device that can compete in cost and usability with tablet pharmaceuticals. The novel design of the Wearable Disposable Electrotherapy platform addresses interdependent constraints spanning automatically initiated and controlled dosing, power density, packaging, and scalable manufacturing with only common, environmentally benign, and nonhazardous materials. The device design supports broad application-specific flexibility (dose, placement) (Fig. 1a).

1. The central innovation of this platform is avoidance of conventional electronics (e.g. printed circuits, heavy metals) which hinder a single-use (environmentally responsible disposable) device. The therapeutic dose is controlled through a printed structure using modular battery cells with interconnects. Together, the 3D battery pack structure, cell shape (tailoring areal energy density to thickness), active materials, and mass inventory provide the requisite voltage and dose control. Dose ramping is achieved through power-load interface design (internal battery pack resistance, electrodes, hydrogel thickness, and ion mobility) based on the progressive impedance changes associated with device application / removal.
2. The common (embedded) substrate for all power/interface components removes any steps by the user (“no assembly required”). Devices are activated upon contact, where the body completes the device discharge circuit (Fig. 1e). Therefore, to use the device, one needs only to apply it (e.g., absence of any controls, even a start button).
3. The platform’s 3D architecture is manufactured entirely using additive/subtractive fabrication with common/benign materials including active materials based on alkaline Zn/MnO_2_ electrochemistry. Moreover, the battery does not require charging; it is fully activated during fabrication.
4. Packing, power (maximum current, capacity), shape, and conformability design requirements are addressed in a novel iterative design workflow (Fig. 2). There is an overlay of electrochemical design, working-temperature regulation (Sup Fig 6), mechanical design to support needed conformability, and sealing structures (including venting system; Supplementary Fig 2d, Supplementary Fig. 3c, Supplementary Fig. 3f).

**Fig. 1.**
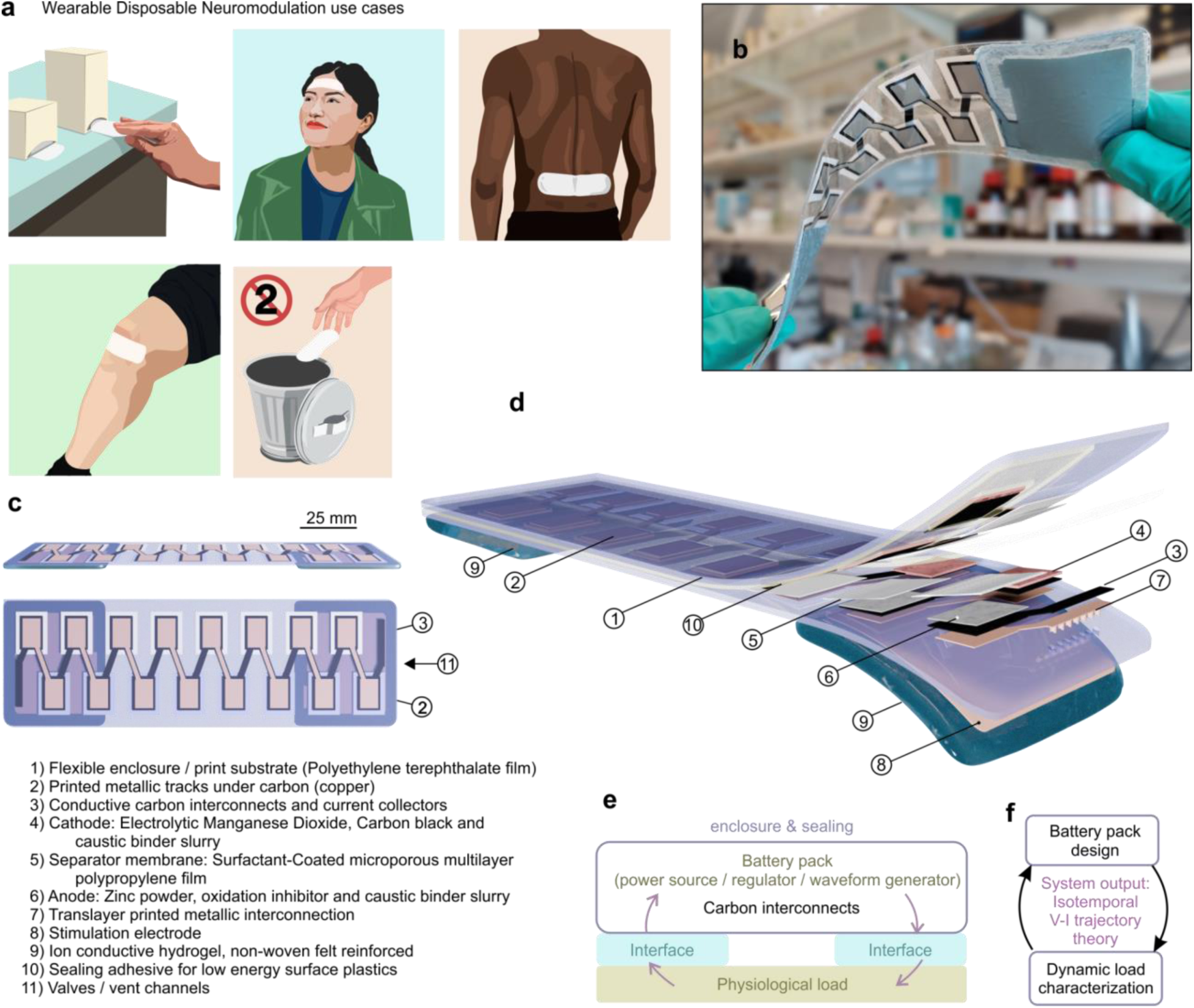
Wearable Disposable Electrotherapy Device. **a)** Application-specific single-use devices are distributed and used akin to disposable bandages or pharmaceuticals. Applications include brain/cranial nerve stimulation (e.g. cognition, headache), electrical stimulation for pain, and accelerated wound healing. Devices are disposable as they are made without conventional electronics, using environmentally-benign materials. **b)** Photograph of exemplary device. **c,d)** Device performance is enabled by a novel layered geometry of active materials and interfaces (element types 1-11) printed onto a common substrate (which becomes the device enclosure). **e)** System diagram: The battery pack discharge is initiated and governed by interaction, through the interface, with the target physiological load. Application-specific therapeutic dose (e.g. neuromodulation, transdermal drug delivery, wound healing bandage) is thus controlled by the device shape and battery pack design. **f)** Unlike prior neuromodulation or battery technologies, discharge is neither current nor voltage controlled. Rather, device chemistry and architecture (battery packs) are designed based on a novel discharge theory.

**Fig. 2.**
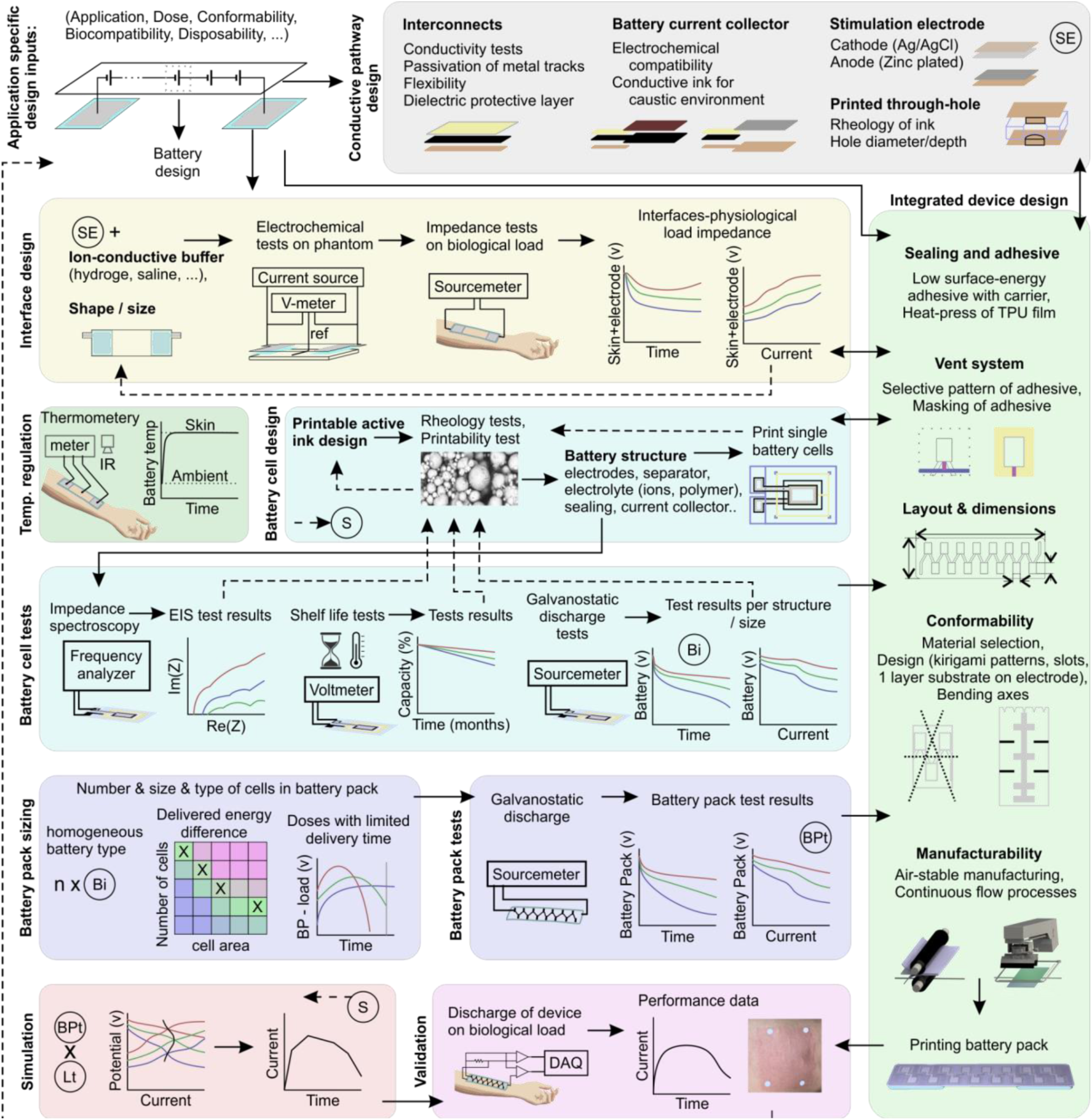
Novel design pipeline enables Wearable Disposable Electrotherapy. A comprehensive workflow and theoretical background support design and validation of application-specific Wearable Electrotherapy Devices. Each design stage (colored regions) incorporates constraints derived from Wearable Disposable Electrotherapy features, design inputs (application-specific dose), and design outputs from other stages and produces design outputs. As the outcomes of each stage impact other stages of the design, device design is integrated and iterative. Design outputs/elements (bold). Integrated workflow (solid arrows). Verification/validation (dashed arrows).

Consequent these features, Wearable Disposable Electrotherapy devices may be distributed and used as economically and simply as pharmaceuticals. Our overall approach is thus orthogonal to efforts creating increasingly complex wearable electronics (9, 14, 15).

Achieving these features involved the innovation of an integrated design procedure (Fig. 2) - demonstrated and validated in detail for an exemplary platform and three electrotherapy applications. Replication of efficacy by Wearable Disposable Electrotherapy requires only the imitation of dose (16), namely current over time and interface shape/position. For each application, dose is, therefore the design input against which device elements (design output) are verified/validated. The design process is iterative (Fig. 2 arrows) but explained below by design outputs (Fig. 2 bolded).

For dose control, the fundamental challenge is the nonlinearity of both energy sources (battery pack) and interfaces-physiological loads - unlike prior electrotherapy devices with electronic output control. A novel design process, governed by application-specific design inputs, separately: 1) characterizes interface-biological impedance loads across subjects; and 2) develops self-regulated battery cells, which are sized into battery packs. Then, the discharge of a designed battery pack through the interface/biological loads is simulated - supporting iterative battery/pack optimization, to ensure reliable output control across subjects. In order to simulate the interaction of the independently characterized (uncoupled) subsystems during operation (coupled), a novel isotemporal-trajectory theory is applied (Supplementary Analysis 1).

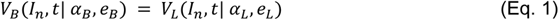

Where the voltages *V*_*B*_ and *V*_*L*_ are the independently characterized battery pack and interfaces-physiological load subsystems, respectively; *I*_*n*_ is a constant current stimulus; *α*_*B*_ and *α*_*L*_ are internal parameters of each system; and *e*_*B*_ and *e*_*L*_ are environmental factors affecting each subsystem. Because battery packs performance can be simulated, human trials are therefore limited to optimized devices.

Interconnect design involves electrically connecting all device elements across a 3D architecture. Although interconnects, through-holes, battery current collectors, and stimulation electrodes are fabricated concurrently they are functionally (electrically / electrochemically) distinct.

Interfaces to the physiological load consist of the stimulation electrode (anode/cathode) and an ion-conductive buffer (hydrogel sheet, non-woven sponge). Interfaces are tested for electrochemical capacity (17; 18) on a phantom, and the design refined accordingly (e.g., printing vs. electrochemical corrosion vs. electroplating), leading to interfaces-physiological impedance load characterization (uncoupled) using a sourcemeter: current levels straddle the application-specific operating range, applied for the targeted duration/charge, with a compliance voltage limit. This selected voltage critically impacts the performance of the final prototype (coupled system), including current ramp-up time and peak, and so battery pack sizing.

Battery cell design is an iterative process involving architecture, anode/cathode inks, current collectors, separator membrane, and electrolyte. Battery cells with different designs and sizes undergo a range of verifications (e.g. galvanostatic discharge, shelf life, EIS). Galvanostatic current levels (range) match those used for interface-physiological impedance load tests.

As single batteries have insufficient potential for delivering dose, a battery pack consisting of a series of batteries is required (Supplementary Fig 1c). Battery packs should contain the required amount of energy and a self-limiting mechanism, with an initial (peak) voltage matched to the compliance voltage limit of the interfaces-physiological load tests. Battery pack sizing includes selection of the number of battery cells, cell types, and size of cells. Battery packs may be homogenous (with a single battery cell type/size) or inhomogeneous - which enhances flexibility in tracking the prescribed dose. Sized battery packs are fabricated and galvanostatically tested using the established current levels (range). Thus the voltage limit and current levels are matched for characterization of the two subsystems: the interfaces-physiological load and battery pack.

Given data from the two uncoupled subsystems (battery pack and interfaces-physiological load) the behavior of the coupled system is simulated using isotemporal-trajectory theory (detailed in Supplementary Analysis 1). The numerical solution provided by this theoretical framework simulates device discharge, which is compared with dose design inputs to consider modifications to the battery pack in an iterative manner.

These system design processes yield application-specific battery pack configurations, developed in parallel with sealing techniques, venting system, comfortability design, all under rigorous limits on acceptable manufacturing methods (scalable, economical, and environmentally benign). Final prototypes are tested against design inputs in human trials to validate each application of disposable Wearable Disposable Electrotherapy.

### Device structure

Disposable Wearable Electrotherapy device elements (Fig. 1c,d, Fig. 7 aii, bii, cii) include the electrochemical cells connected in series (19) (Supplementary Fig. 1c), interconnects and interface elements (stimulation electrodes and hydrogel), sealing and venting system, skin adhesive/dressing based on application. The design must integrate discharge tuned per application, additive manufacturing and robust sealing, and high conformability.

Printed battery cells consistent of an associated anode and cathode, with corresponding cell terminals, are sealed to prevent electrolyte losses as well as to minimize cell-to-cell parasitic losses that result from electrolyte sharing between cells (Supplementary Fig. 2). The anode terminal of the first cell and the cathode terminal of the last cell in series comprise the terminals of the battery pack (power supply; Supplementary Figure 1c). These terminals are connected to the interface, consisting of the stimulation electrode and ion-conductive buffer, which in turn provide connectivity to the body (Fig. 1e).

The batteries adapt primary aqueous alkaline Zinc/Manganese Dioxide chemistry. Electrolytic Manganese Dioxide (EMD) is the cathode active material while metallic zinc is the active anode material (Supplementary Fig. 4). This chemistry offers safety, high energy density, and appropriate self-limited discharge rate (20, 21). The anode and cathode are separated by a specialized membrane made from a porous PP film laminated to a non-woven PP, coated with a hydrophilic surfactant for aqueous applications. The membrane is suitable for extreme pH levels, and it maintains mechanical stability once wet. High device discharge rate is achieved by this high-porosity membrane, high surface area of each cell, low cell thickness, and high conductivity interconnects. As the application is not steady-state (i.e. current changes as a function of time), EIS is a standard tool to characterize the dynamic behavior of varied cell constructs (Supplementary Fig 4 h-j), to then inform application specific battery packs. Hydrogen gas generation (which can deform the enclosure) is lowered using a zinc alloy (with <100 ppm of indium and bismuth), and corrosion inhibitor additive to the alkaline electrolyte and is managed using a vent system (individual battery vents converging to one central channel of 1 mm width, Supplementary Fig. 2d, Supplementary Fig. 3c), which also supports effective vacuum sealing.

In order to fit the required number of cells in series within the physical constraints of the device area and supporting manufacturing, we have developed an innovative cell printing and packing approach. Anodes, cathodes, and their connections are printed in successive steps, in a symmetric geometric fashion on a plastic substrate. These printed sheets form the device enclosure. The interconnects between anodes and cathodes produce the required in-series connection between cells. In the final manufacturing step, the sheet is folded along its symmetry axis, marrying each anode element to its corresponding cathode element (of each cell), resulting in a fully functionalized battery pack (Supplementary Fig. 1a).

The device includes regions of relatively high flexural rigidity (battery cells) bisected by axes of low flexural rigidity (Supplementary Fig. 1bi). This mechanical design, that follows from the planer interlaced battery pack design, supports device conformability. Application specific design may further incorporate other design elements for conformability including articulated enclosures pattern cuts, or regions of low flexural rigidity (e.g. interface regions with a single substrate (Supplementary Fig. 1biv).

The substrate material must support sufficient surface adhesion and compatibility with ink solvents and the extreme pH conditions of battery materials, while also providing the required mechanical characteristics for the application. Materials, such as Polyethylene Terephthalate (PET), are favored for their disposability, cost-effectiveness, and environmentally friendly properties, aligning with the demands of Wearable Disposable Electrotherapy applications.

Interconnects are printed on both sides of the substrate; on the inner side (upon folding) connecting sequential batteries and on the outer side connecting battery pack terminals to the electrode of the interface (Supplementary Fig. 2; Supplementary Fig. 3). The interconnects consist of a narrower copper track beneath a wider carbon conductive track, topped by a broader dielectric layer. This architecture results in interconnects with 75% greater conductivity.

The device is coupled to the skin through the interface elements (hydrogels), facilitating electrical charge delivery and physical support of the device on skin. The interface elements (on the exterior of the substrate) are electrically connected to the battery pack (inside the pack). Though-holes connections are arrays of micrometer-sized holes perforating the bottom substrate (Supplementary Fig. 3b). For the implementation of through-holes, metallic ink is used, and the geometry/size (um range) of the holes through the substrate are adjusted based on the rheology of the metallic ink.

The battery current collector for the anode is printed using copper ink. For the cathode, carbon ink alone or on top of metallic ink is necessary to prevent the corrosion of the current collector in contact with the cathode material. For anode stimulation electrodes, zinc ink, and for cathode stimulation electrodes Ag/AgCl ink printed on a copper surface are options providing higher capacity stimulation electrodes for DC applications.

The exemplary Disposable Wearable Electrotherapy device is designed for low-intensity transdermal stimulation applications (7, 22) with a target dose of 3 ± 1 mA DC average over 20 min (single-use ∼150 mC/cm^2^ capacity) in a conformable packaging with 25 cm² interface electrodes (12 cm inter-electrode distance). A significant design aspect involves ensuring tolerability by limiting current transients at the initiation (**τ** > 1min) and end of stimulation (managed in conventional electrotherapy equipment with microcontrollers) and instantaneous peak current not to exceed 5.5 mA. Hydrogel electrodes are designed for reliable current passage (>0.5 mm thick; volume resistivity of ∼500 Ω.cm; (23), biocompatibility (ISO 10993-5, ISO 10993-10), and provide mechanical adhesion of the device to the skin (moderate skin pull-off adhesion, 20 - 50 g/cm; relative high adhesion on the device side, >100 g/cm), with no residue (e.g. felt reinforced) or irritation.

The exemplary Disposable Wearable Electrotherapy device (Fig. 1b,c,d, Supplementary Fig. 3) comprises of printable layers forming a battery pack (five layers), conductive interconnects (four layers), PET sheets enclosure (two layers), sealant (three layers), polypropylene (PP) strips to mask sealant as normally closed valves connecting inner space of batteries to vent channel, and ion-conductive hydrogels. The 3D design results in a varied number of layers depending on position along device plane - from 3 at enclosure ends, to 9 over internal battery cells, to 11 at conductive hydrogels (not including vents and through holes) (Supplementary Fig. 2b and 3c).

### Scalable manufacture

The manufacturing process for Disposable Wearable Electrotherapy devices adapts methods for economic efficiency and high volume (Fig. 3a). These processes are characterized by subtractive and additive approaches to produce an enabling 3D device structure (Fig. 3b, Supplementary Fig. 1a).

**Fig. 3.**
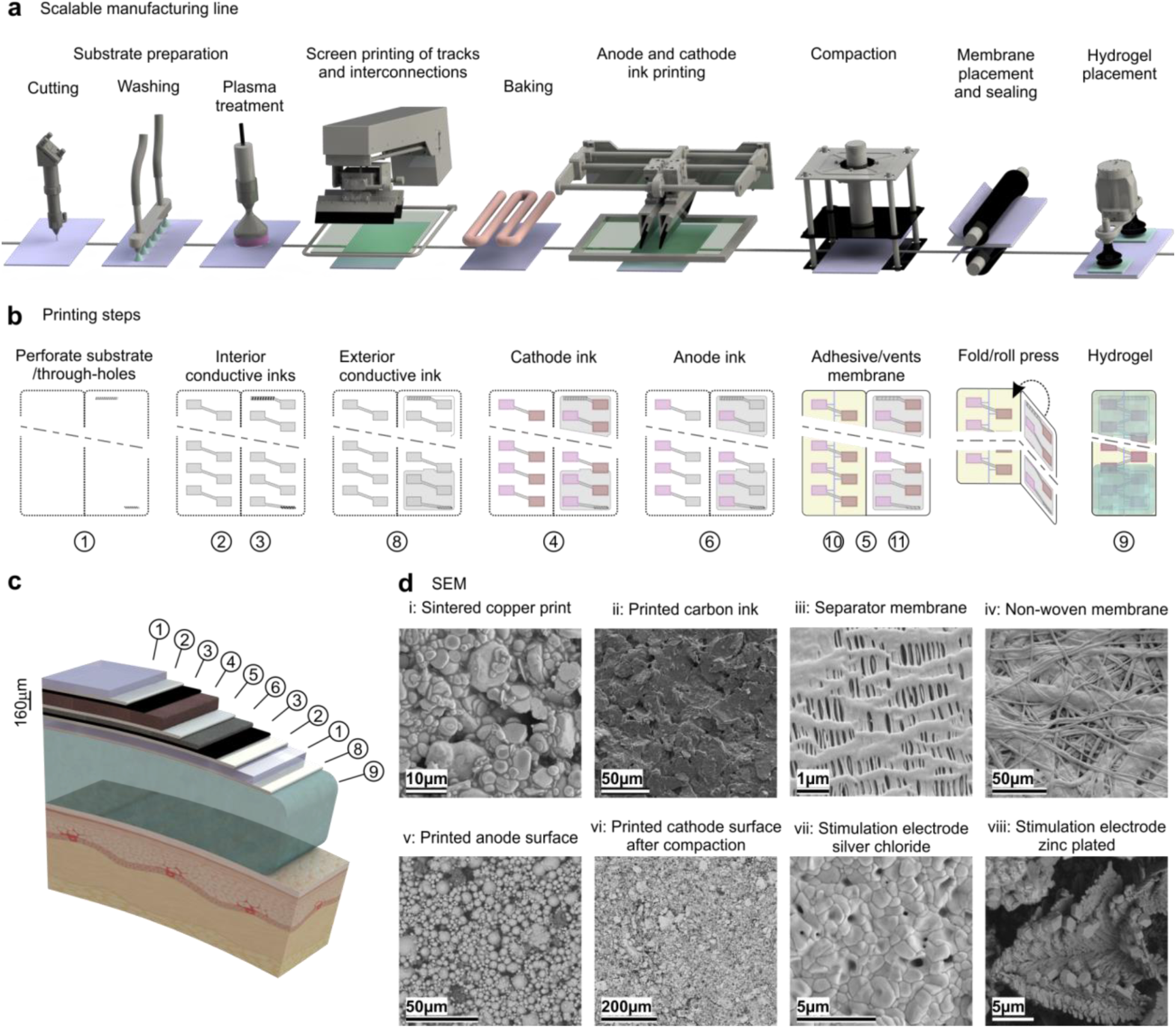
I Wearable Disposable Electrotherapy manufacturing processes. **a**) Scalable manufacturing line consists of substrate preparation, printing conductive/resistive and dielectric inks, curing and baking, printing active materials, compacting, membrane placement and sealing and hydrogel placement. **b**) Steps of additive manufacturing on a single substrate leading to 3D device architecture. **c**) Schematic of layers forming the device including printed battery pack, interconnects, interface, and enclosure elements. **d**) SEM of i: sintered printed copper surface, ii: printed conductive carbon ink, iii and iv: surface of porous PP film and non-woven PP film for retaining battery electrolyte as separator or electrolyte bridge, v: printed anode surface, vi: compacted cathode surface vii: corroded surface of printed silver for cathode stimulation electrode, viii: zinc plated surface for anode stimulation electrode. Elements 1-11 labels as per Fig. 1.

Fabrication begins by heat treating the substrate to ensure its physical stability. This treated substrate serves as the foundation for the deposition of the components that will comprise the electrotherapeutic device. The substrate undergoes rinsing (e.g. isopropyl alcohol) to remove contaminants that can affect the adhesion of the components that will be printed onto the substrate. Plasma treatment modifies the substrate’s surface energy, increasing the polarity of the polymeric substrate, and thus enhancing the ink adhesion and ensuring effective sealing in the downstream processes.

The conductive pathways required by device design are formed by printing conductive and dielectric inks providing required functionalities. Copper ink, conductive carbon ink, zinc ink, Ag/AgCl ink and dielectric ink are among the materials used to construct the various elements of the device. Each fabrication stage is followed by drying and baking steps to fix the features. After forming these features, the battery anode and cathode materials are deposited on the battery current collectors. Electroplating (for microgram accuracy in the deposition of active materials), screen printing (for thinner layers), and/or stencil printing (for controlled deposition thickness, favorable for cathode deposition) were applied for additive manufacture of device layers. The selected inks formulation supports air-stable production, with ink rheology optimized for the printing method while achieving the electrical performance required such as discharge rate. We use a compaction process to increase electrode density that leads to decreased internal battery resistance for the cathode material.

We use a parallel plate battery structure where a separator membrane is assembled between the battery electrodes to electrically isolate them while allowing ionic transport. The sealing stage closes the battery pack between the two enclosure sheets with double-sided tape with an adhesive tailored for low surface energy polymer surfaces. During the sealing stage, the venting system is created by selective masking of adhesive (Supplementary Fig. 2b,d and Supplementary Fig. 3c,f). After sealing, electrolyte is introduced to cells resulting in an active, charged battery unit. In the final fabrication step, hydrogel or non-woven felt (later soaked with electrolyte) as stimulation ion-conductive buffer is applied to the outer surface. Fig. 3c shows the resulting stack of different layers forming the device.

The structure of the battery electrodes and the film uniformity was confirmed by imaging using Scanning Electron Microscopy (SEM) of the deposited layers (Figure 3d). The separator membrane was also imaged using SEM to confirm the absence of defects that could result in short-circuiting a particular cell.

### Interfaces-physiological impedance load characterization

Unlike prior electrical stimulation devices that are current-controlled or voltage-controlled, Disposable Wearable Electrotherapy output is governed by the electrical coupling of the dynamic behavior of the battery pack, the interfaces (electrode, ion-conductive buffer), and the physiological load characteristics (Fig. 1f). For each application (target electrotherapy dose) the interfaces-physiological impedance must be characterized as part of device design (Fig. 2). For this, a device with only the interface components (i.e. devoid of battery pack material) is powered by a sourcemeter (Fig. 4a). The sourcemeter is connected to the stimulating electrodes and programmed to contrast currents (around the target electrotherapy dose) with a voltage-compliance (reflecting the maximum expected from a given battery pack). Because Disposable Wearable Electrotherapy is applied to the body in a pre-energized state, for interfaces-physiological impedance testing the source-meter is activated (to the compliance voltage) prior to placement of the interface components on the body.

**Fig. 4.**
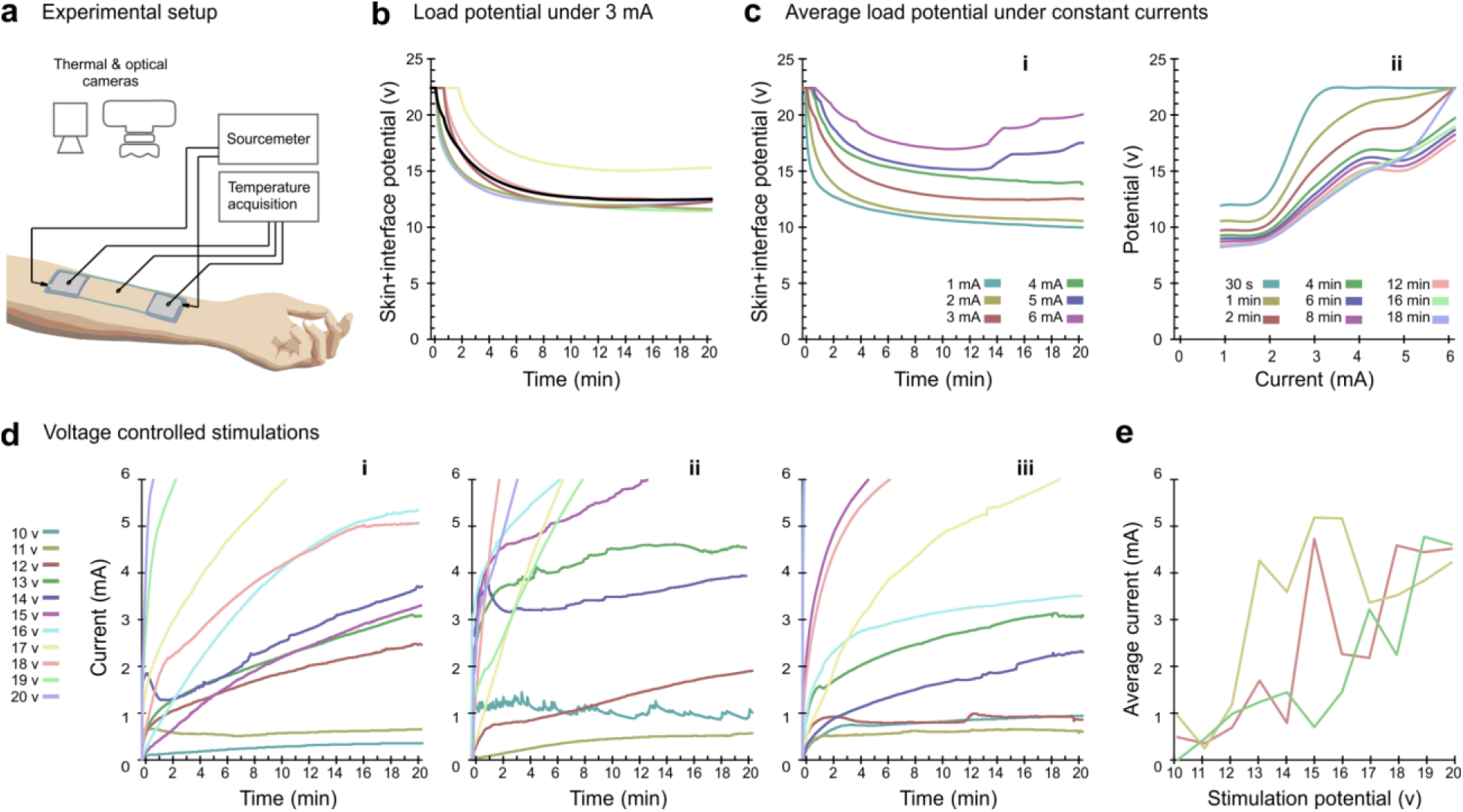
I Load characterization for exemplary Wearable Disposable Electrotherapy device design. **a)** Experimental setup using the interface test device. **b)** Potential of load (skin+interface) under 3 mA over 20 min for individual subjects (colored lines) and average (black line). **c)** i: Average load potential over 20 min under constant currents (used to size battery packs), associated ii: isotemporal V-I curves (used to simulate battery pack discharge). **d)** Voltage-controlled stimulations over 20 min for three subjects (**i, ii, iii**) **e)** average current (over the active duration) for fixed applied voltages. Note unreliable non-monotonic relationship for voltage-controlled stimulation.

To characterize the interfaces-physiological impedance load for the exemplary device, interface-components were applied to subjects’ forearms connected to sourcemeter providing 1-6 mA current-controlled with a 22.4 V compliance. In separate experiments, we considered the response to constant voltage stimulation with 10-20 V in 1 V increments (17 total conditions across n=10 subjects).

Under constant 3 mA stimulation, voltage gradually decreased in each subject (Fig. 4b). On average, across subjects, voltage decreases gradually under 1-4 mA constant currents, but increases at ∼14 min for 5-6 mA current (Fig. 4ci); these relationships are summarized in isotemporal lines (average load, Fig. 4cii; individual subjects’ load, Supplementary Fig. 5) for subsequent battery pack design. The interfaces-physiological impedance is dynamic as a function of applied current and time, reflecting nonlinear processes at the interface electrode (17) and skin (24, 25).

Batteries do not provide constant current and their internal voltage/impedance is a nonlinear function of the current drawn; this creates a complex interdependence between source and interfaces-physiological impedance load. In subsequent design steps (Fig. 2), we show how interfaces-physiological impedance data informs novel battery pack design that accounts for (and indeed leverages as a self-regulating mechanism of battery voltage; Supplementary Analysis 1) a dynamic coupling between batteries and interfaces-physiological impedance load.

Constant voltage stimulation (10-20 V) produces unreliable current (Fig. 4d). Current fluctuates on an experiment-wise basis over time. Current is not monotonic with time or applied voltage. These results confirm that voltage-controlled stimulation (i.e. from an idealized battery) is not reliable for stimulation across interface and physiological impedance loads. Our specialized battery packs are designed to change in voltage with interfaces-physiological impedance load.

### Battery pack design

For the exemplary specific target dose (3±1 mA, 20 min) and associated interfaces-physiological load dynamics, a homogeneous battery pack (power source with limited voltage and current) is designed. Provided single battery cell electrochemistry parameters (ink formulation, screen thickness, separator type, electrolyte composition) and device structure (3D connectivity, fabrication/sealing technique) designed in prior stages, we next address adjusting two dimensions to optimize battery pack performance: cell size and number of cells. Notwithstanding our illustration of the exemplary Disposable Wearable Electrotherapy device design in sequenced stages, we emphasize system design (Fig. 2) is iterative.

We conducted an array of single battery cell discharge tests, maintaining a consistent discharge rate of 3 mA over 30 min (Fig. 5a). During these tests, cell sizes ranged from 0.8 cm^2^ to 2.4 cm^2^, with constant areal densities of the active materials. We scaled these recorded voltages by factors ranging from 10 to 18 to predict associated projected battery pack voltages under a 3 mA discharge rate. These values are compared with potentials across the interfaces-physiological load under 3 mA for 20 min stimulation (Fig 4). Our objective was to identify a battery size and the required number of cells that minimized the voltage variation throughout the 20 min span (Fig. 5bi) while also ensuring the battery pack neither depleted prematurely nor persisted substantially beyond the 20 min (Fig. 5bii, Supplementary Fig. 4e). At this stage, the battery pack sizing (optimized by cell types, sizes and number) was based on minimizing the energy difference, over the targeted dose, between the battery pack and load by:

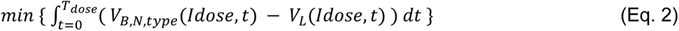

with respect to battery size and time for each of N possible battery cell choices, and

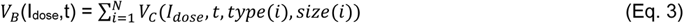

where V_c i_ (I_dose_,t, *type*(*i*), *size*(*i*)) is the voltage of a single battery cell at time t for a given current-controlled discharge (I_dose_) applied over time (T_dose_), which are set to the target dose. The battery type and size of each battery cell affects its voltage, and hence cumulative battery pack voltage. For the exemplary device, the selection of battery chemistry and a homogenous pack design reduces the optimization to cell area and number. Following this analysis, a configuration of 1.5 cm^2^ with 14 cells per pack was deemed optimal for the specified target dose.

**Fig. 5.**
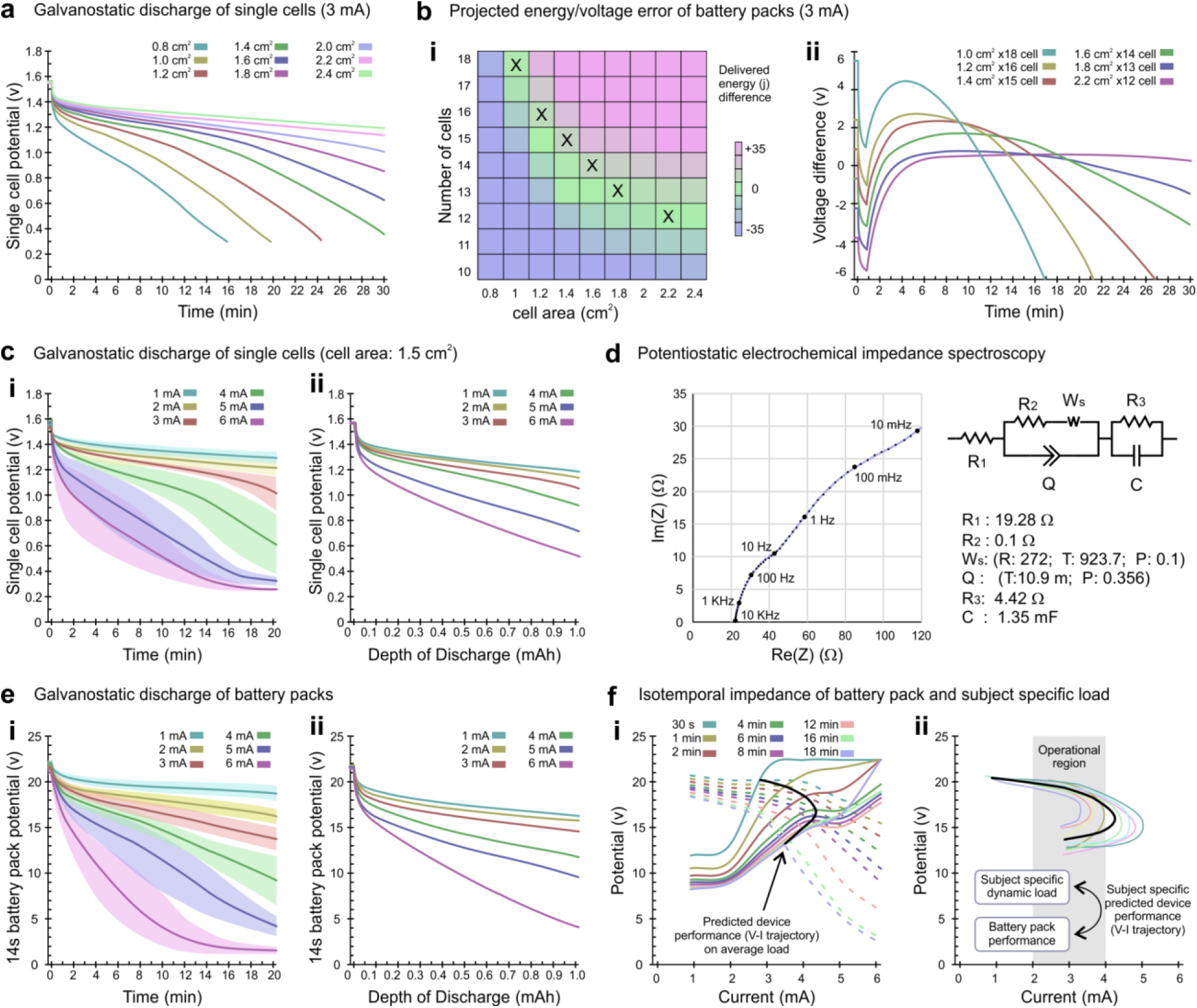
Battery cell and pack design (sizing), validation, and stimulation for exemplary Wearable Disposable Electrotherapy. **a)** Galvanostatic discharge curve of single-cell batteries (average) with different cell sizes under 3 mA current over 30 min; **b i).** Projected energy (j) error of battery packs with 10 to 18 cells made from cells with different sizes under 3 mA discharge vs average of energy required to stimulate load with 3 mA during 20 min (x: selected designs for next stage); **b ii)** Voltage difference between six battery pack designs (selected based on minimal projected energy error) and voltage of load over 30 min at 3 mA. **c)** Galvanostatic discharge curve of single cells with sized area of 1.5 cm^2^; **d)** Potentiostatic electrochemical impedance spectroscopy of single cell per 1 cm^2^ area over frequency range of 10 mHz to 10 kHz with bias equal to OCV of freshly assembled battery and fitted model; **e)** Galvanostatic discharge curve of sized (14 cell, 1.5 cm^2^ area) battery pack, **i**: over 20 min time, **ii**: over-discharge depth; **f i)** Isotemporal V-I curve of sized battery pack discharge (dashed lines) with overlaid average isotemporal V-I load curve (solid line). The intersection of these lines is the predicted discharge for the sized device into the average load (black line). **f ii)** Given each subject’s V-I load curve, a subject-specific (colored lines) and average (black line) sized device discharge is simulated.

We then measured (n = 72) the average potential of single 1.5 cm^2^ cells over a 20 min span across various discharge rates (1-6 mA); both average performance (Fig. 5ci, solid line) and performance variation due to variation in battery fabrication (Fig. 5ci, shaded). The average potential of the cells for up to a discharge depth of 1 mAh was calculated (Fig. 5cii).

We further characterized the performance of single battery cells using potentiostatic electrochemical impedance spectroscopy (EIS) and developed an associated circuit analog model (Fig 5d). The lumped-parameter device impedance model reflects electrochemical processes occurring in the cell (26). These single cell data are then used to inform iterations of battery chemistry (Supplementary Fig 4 h-j) and battery pack design.

### Verification and system simulation

Having designed the size and quantity of batteries for each pack for the exemplary target dose and load, we fabricated and subjected these battery packs to galvanostatic discharge tests (n= 60 battery packs). The average potential of battery packs over a 20 min span across various discharge rates (1-6 mA) was measured; both average performance (Fig. 5ei, solid line) and performance variation due to variation in battery fabrication (Fig. 5ei, shaded) is measured. The average potential of the battery packs for up to a discharge depth of 1 mAh was calculated (Fig. 5eii).

The battery pack discharge dynamics is represented in isotemporal lines (colored dashed lines; Fig. 5fi). These are overlaid with isotemporal lines from the load average (colored solid line; Fig. 5fi). For each time point, the interaction of these lines is represented (solid black line; Fig 5fi). The temporal evolution (over the stimulation duration) of the intersections between the voltage-current characteristics of the battery and of the load is the projected device output. This corresponds to the simulated Disposable Wearable Electrotherapy dosing for an average associated interfaces-physiological impedance load. The projected output is then analyzed across multiple subjects, based on their respective load analysis (Supplementary Fig. 5; Fig 5fii). These trajectories show the projected operational envelope of the device (within a safe range). Reflecting our device design (sizing and battery cell chemistry; Supplementary Fig. 4g), trajectories show the current will initially ramp up with limited decrease in voltage. As stimulation proceeds, the current reaches a maximum value, after which the voltage decreases substantially, limiting current delivery. A further decrease in voltage ramps the current down.

As the device is thin and battery pack (polymer) elements are impacted by temperature, we quantified device operational temperature. Sensors embedded in the device (on the anode stimulation electrode, device center, and cathode stimulation electrode) showed devices quickly (**τ** = 40 s) warm from room temperature to surface skin temperature (Supplementary Fig. 6a), with an incrementally higher temperature at the cathode (Supplementary Fig. 6c, consistent with enhanced erythema). Associated warming at the skin surface was confirmed by thermal imaging Supplementary Fig. 6d. We further developed a computational model of device heat transfer (Supplementary Fig. 6b), consistent with experimental findings. These results confirm that primary temperature increase reflects device warming to body surface temperature, consistent with conventional electrotherapies (27), during the initial (ramp up) discharge period.

Stability (shelf-life) testing of the exemplary Disposable Wearable Electrotherapy pack assessed the battery pack’s capacity. Over 15 days, the average voltage decreased by 5.25 mV per cell daily. Allowing for a ∼20% decrease (e.g., from 1.55 V to 1.2 V per cell), the results indicate a stability of 66.7 days (i.e., over two months of stability at the prototype stage).

### Validation

With chemistry and sizing designed for the exemplary application, the exemplary Disposable Wearable Electrotherapy discharge performance was validated (n=10 subjects; Fig. 6a). Devices satisfied all other requirements, with a final thickness of 1.25 ± 0.07 mm and weight of 10.5 ± 0.1 g. Devices were applied to the skin (t = 0) for 20 min, and the generated voltage (Fig. 6b) and current (Fig. 6c) measured. Across subjects voltage gradually decreased while current gradually increased (**τ** = 8.7 ± 3.4 min) to a peak (3.6 ± 0.8 mA) while current was sustained. A further decrease in voltage reduces current. Stimulation was sustained for the 20-minute application (average voltage 15.4 ± 2.4 V; average current 2.8 ± 0.7 mA; average power 41.8 ± 12 mW). As the device was removed, the current was progressively aborted (**τ** = 5.0 ± 0.5 s).

**Fig 6.**
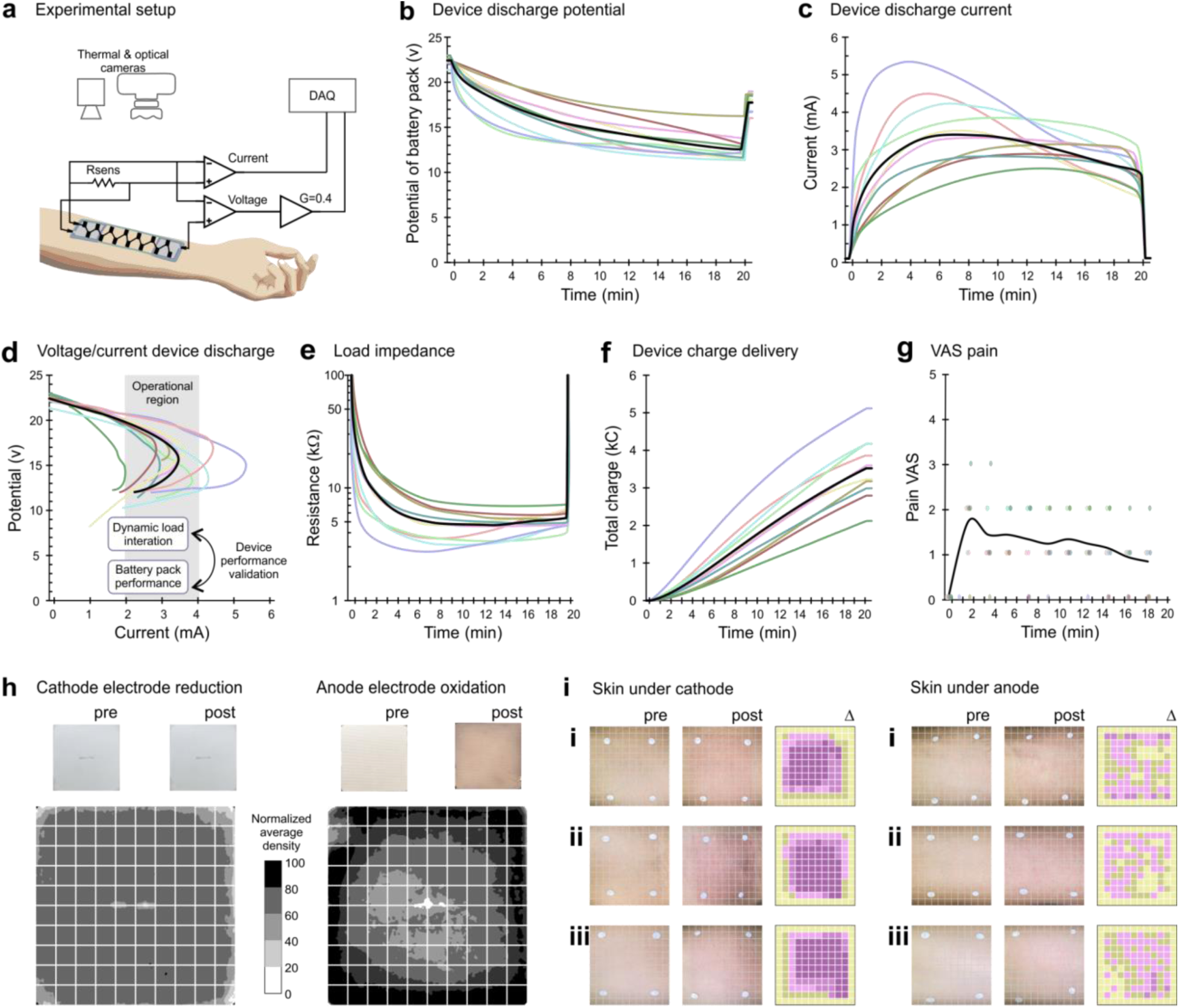
Validation of exemplary Wearable Disposable Electrotherapy self-limited discharge. **a)** Experimental setup for the target application across subjects’ forearms. Individual subjects (colored lines) and average (black) lines are shown. **b)** Voltage of battery packs and **c)** output current over 20 min of stimulation. **d)** V-I curve of device throughout the stimulation. Compare these measured discharges with device-design simulation (Fig 5fii). **e)** Impedance of the load (electrodes + skin), **f)** cumulative charge delivered across the load, and **g)** pain levels over the 20 min of stimulation. **h)** Distribution of stimulation cathode electrode reduction and anode electrode oxidation. **i)** Skin redness relative heat map under stimulation cathode electrode and under anode electrode, for three subjects (**i, ii, iii**).

Voltage-current trajectories (Fig. 6d) exemplify Disposable Wearable Electrotherapy dose control and performance along system-design simulations (compare with Fig. 5fii). Discharge is neither strictly current-controlled nor voltage-controlled, but each subject is governed by the battery pack design given the dynamic impedance response (Supplementary Analysis 1). Load impedance initially decreased with current application (Fig. 6e) but reliably plateaued and normalized across subjects (range 4-7 kΩ at 18 min).

Discomfort during electrotherapy was assessed by conventional VAS rating (23) at 11 time points during stimulation. Stimulation was well tolerated (average pain VAS 1.1 ± 0.75; across 110 measurements never >3; Fig. 6g), with expected transient erythema and no lasting skin irritation. Subject’s transient and mild perceptions of current flow (e.g. “tingling”) are consistent with functional electrical delivery (i.e. activation of nerves; 28).

Uniformity of current delivery was assessed across electrodes and skin surfaces. Charge uniformity was imaged (pre/post discharge, 2D optical scan) at the anode stimulation electrode, as evidenced by oxidation, and at the cathode electrode, as evidenced by gas evolution (Fig. 6h). Average normalized charge density was uniform at the cathode and moderately higher at electrode edges (annular) at the anode. Stimulation uniformity at the skin was assessed by high-resolution photography (pre/post discharge) of skin erythema (Fig. 6i; 29). Immediately post-discharge, erythema was relatively uniform under the cathode while milder and punctate under the anode. At appropriate doses, erythema is expected and transient (29), non-hazardous and consistent with skin stimulation, and - in the context of iontophoretic drug delivery and wound healing - desired (5, 28).

Functional conformability (23) is validated through the combination of impedance stability (Fig. 6e), current density uniformity (Fig. 6h,i), and tolerability (Fig. 6g). By achieving an electronics-absent design with thickness of battery pack (all layers) ranging 400 - 700 µm thickness (Supplementary Fig. 2b), flexibility is governed by the design/layout of the battery pack. Given PET flexural modulus (1.30 - 4.69 GPa), sheet thickness (100 µm), and device to skin surface ratio <0.3, a >97% comfortability is expected (30).

### Applications of Wearable Disposable Electrotherapy platform

We demonstrate three applications of Wearable Disposable Electrotherapy using the directed design framework (Fig. 2). For each application, the electrotherapy dose and mechanical consideration of application serves as the design input to the Wearable Disposable Electrotherapy analog and the validation of design outputs. The range of performance (design inputs) of these applications and exemplary device are selected to demonstrate the platform’s flexibility to broad electrotherapy applications. Current spans 2 orders of magnitude (30 uA to 3 mA), duration (∼20 min to >2 hours) with 0.5-2 cm² battery packs of 4-14 cells. Both homogeneous and inhomogeneous battery pack sizing are demonstrated. Application-specific interfaces span 4.5 to 25 cm² contacts, hydrogel or nonwoven sponge ion conductors, and varied stimulation anode/cathode materials (copper, zinc, silver, carbon and silver chloride). Battery packs and interface are packaged into application specific enclosure (18 to 132 cm²), with mechanical design supporting comfortability, Variations in interfaces and target dose, result in different interfaces-physical impedance loads - reflecting nonlinear dynamics of the load - which are accounted for isotemporal-trejector theory (Supplementary Analysis 1).

Transcranial Direct Current Stimulation (tDCS) is an non-invasive brain stimulation technique (31), trialed for a range of neurological and psychiatric disorders (3,4). A typical dose is ‘bi-frontal’ 1-4 mA (with 30 second ramp up/down), 20-30 min, ∼25 cm^2^ electrode on the EEG 10-10 F3/ F4 scalp positions. Bi-frontal Wearable Disposable Electrotherapy design inputs were with electrodes below hairline, using 3 ± 1 mA DC average to provide target electric field to the frontal cortex (max 2.4 V/m at the dorsolateral prefrontal cortex; Fig. 7a; 32) with ramp up **τ** >30 s and instantaneous peak current <4 mA. This design was based on interfaces-physiological impedance load under constant currents (1.5-4 mA; Fig. 7aiv). Integrated system design (Fig. 2) produced an articulated device architecture (Fig. 7ai) including 4 inhomogenous cells (Fig. 7aii). Conformability is enhanced through the use of a single substrate over the interface (Supplementary Fig. 1biv) and bend-line cuts in the substrate, which facilitate deformation to the forehead (Supplementary Fig. 1bii). To deliver the prescribed dose, battery pack sizing (eq. 2) resulted in an inhomogeneous configuration with 3 cells (1.8 cm^2^) with 34 wt% KOH electrolyte and 1 cell (1.8 cm^2^) with 0.7 wt% PAA polymer added to electrolyte that limits the peak current (Supplementary Fig. 4g). Designed devices were prototyped according to our manufacturing process (Fig. 3) and validated on human subjects (Fig. 7avii) demonstrating discharge performance within design specification and matching design theory.

**Fig 7.**
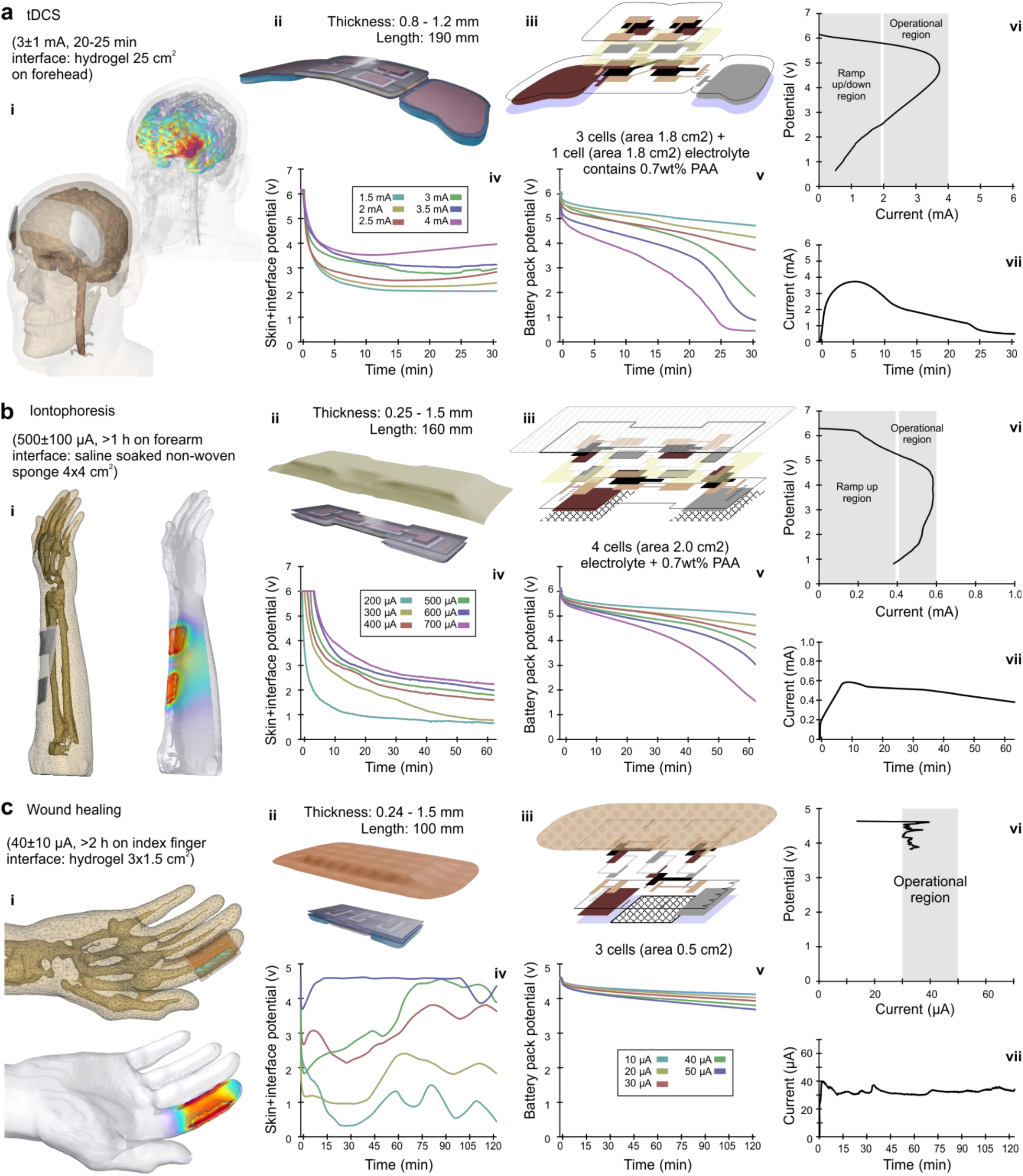
Application specific Wearable Disposable Electrotherapy design. The design pipeline validated for the exemplary device is applied to three use-cases: tDCS (a), iontophoresis (b) and accelerated wound healing (c) - selected for efficacy supported by dozens of RCTs such that a Wearable Disposable Electrotherapy needs to reproduce the dose (design input). **i)** For each application high-resolution MRI-derived models simulate the desired tissue current delivery. **ii)** The outcomes of an integrated design process are application specific Wearable Disposable Electrotherapy devices, Thicknesses scaled 10x for clarity. **iii)** Exploded view of layered geometry of active materials and interfaces. **iv)** These designs are the outcome of a process including application dose-specific interfaces-physiological load impedance measurements, informing battery cell design and battery pack sizing. **v)** Galvanostatic battery pack prototype discharge informs **vi)** isotemporal device simulation. **vii)** Application of assembled devices to subjects produces successful outputs.

Iontophoresis is an established therapy passing DC through the skin for applications including hyperhidrosis and transdermal drug delivery (33). A conventional iontophoretic dose uses ion-carrier non-woven sponge interface (4×4 cm) with charge rated dose >30 mA-min (∼1.8 C); when applied for 60 min sustaining an average current ∼500 uA (34). These served as the design inputs for the iontophoresis application Wearable Disposable Electrotherapy (Fig. 7b). System design (Fig. 2) yielded a Wearable Disposable Electrotherapy iontophoresis design (Fig. 7bii) using 4 homogeneous cells (2.0 cm2) with 0.7 wt% PAA polymer-supplemented electrolyte to self-limit the peak current. The conformability was enhanced by narrower substrate between two interfaces (Supplementary Fig. 1biii). Current flow simulation predicts resulting charge density of 31.2 µA/cm2 per second at the device-skin interface (Fig. 7bi). According to isotemporal trajectory theory (Supplementary Analysis 1), the design was developed by measuring the interfaces-physiological impedance load under constant currents (200-700 µA; Fig. 7biv). The battery pack was accordingly sized and discharged (Fig. 7bv) and the iontophoresis-application Wearable Disposable Electrotherapy device built (Fig. 3). Using a skin phantom, enhanced diffusion of ionic dye (molecular weight similar to the range of common drugs used in iontophoresis) was verified (Supplementary Fig. 7). Finally, devices were validated in human trials to produce the prescribed discharge performance (average current 472 µA, 32.7 mA-min in 60 min; Fig. 7bvii)

Electrical stimulation can accelerate wound healing (5,6,35) Effective doses include low-intensity DC at 30-50 µA average over >2 hours. An integrated design process (Fig. 2) resulted in a wound healing-application Wearable Disposable Electrotherapy design including three battery cells, and stimulation electrode hydrogels on both sides of a wound dressing pad. The manufactured device is then placed on a skin adhesive for bandages. The substrate is made from a stretchable thermoplastic film with slots between each battery cell for additional flexibility (Fig. 7cii, Supplementary Fig. 1bii). During heat press sealing, the stretchable thermoplastic film undergoes copolymerization, eliminating the need for a separate adhesive layer between the two substrates. Using an interface device and sourcemeter (10-50 µA), the interfaces-physiological impedance load was determined. Battery packs were sized according to isotemporal trajectory theory (Supplementary Analysis 1) and the device was built accordingly. Current flow simulation predicted a resulting largely uniform current density through the targeted region (Fig. 7cii). Wearable Disposable Electrotherapy wound healing devices were then validated (Fig. 7cvii) demonstrating discharge performance within design inputs and matching design theory.

### Usability, Economics, Environmental, Healthcare Equity Impact

Usability is a barrier to electrotherapy adoption, compliance, and effectiveness (36; 11). Wearable Disposable Electrotherapy devices is a breakthrough from all prior electrotherapy equipment, which require patients to be tethered to an electronic stimulator with numerous steps at each use and between each use. These obstacles complicate the treatment experience, throttle adoption and impair compliance. Wearable Disposable Electrotherapy is a categorical change in usability, for the first-time making electrotherapy use akin to pharmacotherapy or topical medication. By fundamentally simplifying dissemination, storage, use, and disposal (Fig. 8), Wearable Disposable Electrotherapy is a new medical delivery platform. Wearable Disposable Electrotherapy are thin, discreet, comfortable strips, with an application-specific single dose automatically delivered upon application to the skin (Fig. 6, 7). Through our invention of an electronics-free electrochemical architecture, categorically enhanced usability is achieved alongside economical advantages, environmental superiority, and equitable access compared to conventional electrotherapies.

**Fig. 8.**
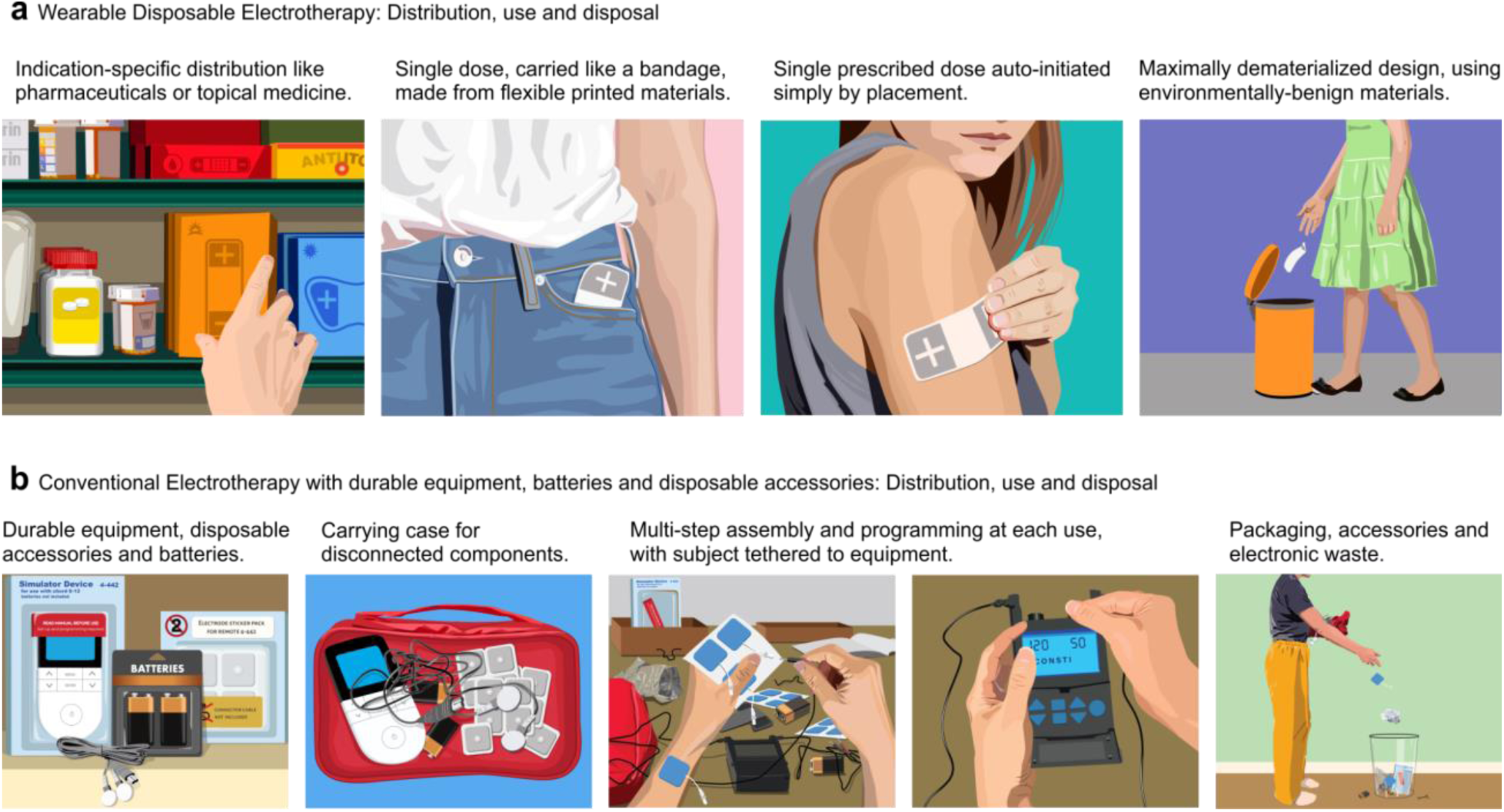
Wearable Disposable Electrotherapy eliminates barriers to use of traditional electrotherapy, and facilitates distribution akin to pharmacotherapy or topical medicine. (Left to right) Distribution: Wearable Disposable Electrotherapy is dispensed in application-specific single-dose strips, similar to distribution of drugs or topical medicine. In contrast, conventional electrotherapy involves both durable and disposable components, as well as power sources. Carrying: Each Wearable Disposable Electrotherapy, containing a single dose, is like a bandage, while transporting conventional electrotherapy requires all components. Application: To use Wearable Disposable Electrotherapy, the strip is simply applied to skin – the device is discrete and automatically initiates and provides a single dose. With conventional electrotherapy a multi-step process involves tethering the electronic stimulator to the patient, using disposable electrodes, and programming/initiating therapy. Storage/disposal: The Wearable Disposable Electrotherapy minimizes environmental impact. Conventional electrotherapy devices have both durable electronics which must be stored and charged (e.g. batteries) between uses (and whose eventual disposal includes toxic materials) as well as disposal of single use electrodes (which in themselves have more metal than a Wearable Disposable Electrotherapy).

Wearable Disposable Electrotherapy economic efficiency is superior to traditional electrotherapy, which includes both durable electronics and consumables (batteries, disposable electrodes). Wearable Disposable Electrotherapy is by design dematerialized to essentials for delivering a single dose (i.e. minute quantities of functional and electrochemically active materials besides materials needed for interfaces; Supplementary Table 1). At the end of a treatment session, device materials are exhausted. In contrast, traditional devices: 1) Consume energy inefficiently (e.g. displays, voltage step-up) using stand-alone batteries (with inevitable waste); 2) Entail significant costs for materials, manufacturing (with higher Product Complexity Index), packaging, and shipment, amassing hundreds of grams of molded plastics, PCBs, electronics, and connections; moreover 3) This burden of durable equipment still requires disposable electrodes (∼10 grams per use including metal connectors; Supplementary Table 1). Wearable Disposable Electrotherapy prioritizes scalable manufacturing processes that are air-stable and less energy-intensive, unlike costly material and assembly processes for electronic stimulators.

Studies established traditional electrotherapy as cost-effective healthcare, without discounting up-front equipment costs (35; 37). Wearable Disposable Electrotherapy maintains the healthcare quality benefits without upfront equipment or training costs.

The production of traditional electrotherapy devices (electronics) hinges on the utilization of rare earth elements and heavy metals, alongside manufacturing processes that are resource-intensive and detrimental to the environment (toxins and carbon emissions). The end-of-life disposal of traditional devices further compounds their environmental footprint. Wearable Disposable Electrotherapy production and use is inherently sustainable, being dematerialized to essential components, and using only abundant and environmentally benign materials (Supplementary Table 1). Moreover, this unique form is coupled by design (Fig. 2) to scalable (additive) manufacturing techniques that do not depend on toxins, thus minimizing the environmental impact of Wearable Disposable Electrotherapy. The limited usage (inevitable disposability) exacerbates the environmental impact of traditional devices.

Technology-centered health care advances (eg “smart” devices) often preferentially benefit users of privileged socioeconomic backgrounds. Wearable Disposable Electrotherapy decreases healthcare inequity. Conventional electrotherapies have a high startup cost (all durable equipment; 38), while Wearable Disposable Electrotherapy can be trialed with a single disposable device. The multi-step setup, programming, and maintenance of traditional electrotherapy is an accessibility barrier, while the auto-initiated bandage operation of Wearable Disposable Electrotherapy is intuitive (39) enhancing access to broader demographics. Deployability is a third factor for equitable access of medical devices, with Wearable Disposable Electrotherapy not requiring batteries/charging and can be simply distributed.

## Conclusions

In summary, we designed, fabricated, and validated the first electrotherapy platform using additive manufacturing with common environmentally-benign materials - without electronics. Device packing, power, shape, and conformability design requirements are addressed using this design workflow. Electrotherapy dose is “built-into’’ device architecture/chemistry, with discharge initiated simply by applying the device. We develop an associated theoretical framework for device design based on coupling between load and battery pack. Platform manufacture can be scaled and customized with printable circuit technologies (40), non-linear conductive elements for pulse synthesis (OFET; 9), and printable sensor-like materials (e.g., sensitive to temperature, humidity, and pressure).

Wearable Disposable Electrotherapy can be distributed like drugs or topical creams (Fig. 8), as dose/indication specific adhesive strips (Fig 1a). A patient can carry a single device, simply apply it discreetly when needed, and then discard the device. Based on iontophoretic drug delivery, drug alluding strips are thus made practical. The established principle of electrical wound healing can be made practical with disposable electrically driven bandages. Strips applied to the head can provide neuro-psychiatric therapies based on cranial nerve stimulation or cortex stimulation. This paper serves to show these and other applications are technically feasible.

## Methods

### Manufacture

#### Substrate preparation

The substrate is fabricated from 100 µm-thick PET sheets (McMaster #8567K44) which undergo a series of preparation steps: 1) Heat treating of the substrate for the purpose of mechanical stability by baking at 90°C for 15 min; 2) Laser cutting to perforate boundaries of the device and folding line as well as throughholes. This allows the pouch to be a part of the substrate sheet during printing steps, and subsequently to be separated for sealing of the battery pouch. The fold line facilitates accurate alignment of battery electrodes during folding while preventing easy tearing; 3) Washing with isopropyl alcohol followed by air drying; 4) Surface treatment with cold plasma (Relyon plasma piezo brush PZ3) to improve adhesion.

#### Printing of conductive tracks (interconnect, through holes, stimulation electrodes, current collectors)

A copper conductive ink (copprint LF-350) or silver conductive ink (Saral Silver 700) is screen printed (Novastar SPR-45 stencil/screen printer) onto both sides of the substrate (Supplementary Fig. 2a and 2c). Following printing, the substrate is heated in an oven (90C) to facilitate the evaporation of the solvents present in the ink used. After baking, the thickness of the printed silver track is ∼20 µm. Subsequently, a carbon passivation layer (Saral Carbon 700A) is screen printed and baked in a similar fashion; this was only on the inner side with a screen template 1 mm wider than that used for silver from each side. After baking, the thickness of the printed carbon track is ∼40 µm.

To establish an electrical connection between both sides of the substrate, we implement laser-cutting to create micrometer-sized holes in the form of an array in the substrate before printing. During the screen printing of the conductive inks on both sides of the substrate, the ink permeates these holes, creating a conductive path between the two sides of the substrate. The size and number of these holes are tailored to specific ink properties (rheology and particle size) to achieve reliable adhesion and conductivity. To support device verification and validation, three conductive tabs were added to the design at the battery pack terminals and at the cathode stimulation electrode, which allow monitoring of current and voltage.

#### Preparation of cathode ink, anode ink and electrolyte

The cathode ink used for exemplary device is composed of 70 wt% electrolytic manganese dioxide (EMD), 3.5 wt% carbon black as a conductive additive, 5 wt% KOH, 20 wt% deoxygenated DDI water, and 1.5 wt% PVA (average MW 94k) as a binder. The slurry is prepared by mixing water and PVA and KOH in a nitrogen box to prevent oxygen dissolution in water during mixing. This solution then is sealed and kept in the refrigerator at 4 °c. Prior to use, the mix of EMD powder and carbon black powder is added and mixed and finally loaded into a syringe for application. The cathode ink for applications is composed of 79.5 wt% electrolytic manganese dioxide (EMD), 3.5 wt% carbon black as a conductive additive, 15.5 wt% deoxygenated DDI water, and 1.5 wt% SBR as a binder.

The anode ink used for exemplary device consists of 75 wt% zinc powder, 0.4 wt% zinc oxide as corrosion inhibitor, 5 wt% KOH, 17.8 wt% deoxygenated DDI water, 0.25 wt% PAA (average MW 450k), and 1.55 wt% Na-CMC (average MW 90k) as binders. The slurry is prepared by mixing all ingredients, except for zinc powder, in a nitrogen box to prevent oxygen dissolution. The slurry is then sealed and refrigerated at 4 °c. Prior to use, zinc powder is added to the solution inside of a syringe and mixed to create the anode ink. The anode ink for applications is composed of 74 wt% zinc powder, 0.3 wt% zinc oxide as corrosion inhibitor, 23.5 wt% deoxygenated DDI water, 1.4 wt% Na-CMC as filler, and 0.8 wt% SBR as binder.

The electrolyte is formulated with 66 wt% deoxygenated DDI water and 34 wt% KOH.

#### Sealing method

The device is sealed using a double-sided acrylic adhesive designed for low surface energy plastics, with an interior carrier film for enhanced mechanical stability with a thickness (0.17 mm) less than the battery pack cells and interconnects (3M 9495LE). The double-sided adhesive sheet is cut (prior to removing the liner of both sides) using a laser cutter into two strips, each with openings corresponding to cells on one row of the battery pack (Supplementary Fig. 3f). When placed on the substrate a gap between the two adhesive strips forms the central channel of the venting system (Supplementary Fig. 2d and 3f).

#### Placement of separator

The separator membrane (Celgard 5550) used consists of polypropylene film laminated to a polypropylene nonwoven fabric, coated with hydrophilic surfactant for aqueous applications. This membrane has a thickness of 110 µm and 55% porosity, for high electrolyte retention and ion conductivity for high discharge rate. The membrane is laser cut to appropriate size and placed on the cell using cut double sided tape.

#### Vent channel and valves

After placing the membranes on double sided tape, thin strips are printed on the adhesive using non-stick ink (1 mm width), connecting the middle of the battery cell to the vent channel (Supplementary Fig. 2d and 3c). These thin strips mask the adhesive, creating normally closed valves, they provide an escape for air (during battery pack sealing) or generated hydrogen by the cells, to the vent channel.

#### Printing of active materials on current collectors

In the exemplary device each membrane is saturated with 9.5 mg (8 µL) of electrolyte before printing active materials. The volume of electrolyte is crucial to control since insufficient electrolyte reduces battery performance, while excess electrolyte wets the surface of double-sided tape resulting in poor sealing. Then anode and cathode inks are deposited on printed substrates using screen / stencil. Immediately after printing anode ink, double sided tapes with soaked membranes are placed on two anode rows of the device. This prevents printed zinc from drying. By folding the device on its fold line, both sides of the substrate meet in alignment to form a sealed battery pack (Supplementary Fig. 2a). The sealed battery pack goes through a roller from each side to the middle of the device where the vent channel is to push trapped air out of cells through valves.

#### Battery pack quality control

After compressing the pack, terminals of the pack are connected to a multimeter to read the initial open circuit voltage (IOCV). Some manufacturing problems can be detected by observing subtle changes of voltage.

#### Interface hydrogel application

Ion conductive hydrogels (Axelgaard AG625) are cut to the size (45 mm x 56 mm, 25.2 cm^2^) and placed on the stimulation electrodes. As needed, fastening (non-conductive) hydrogel (Axelgaard AG535) can be used in between stimulation electrodes. The hydrogels are covered with PET liner until use. The hydrogel placement step was omitted for battery pack discharge verification tests. The optimization of hydrogel (and associated stimulation electrode material) for the exemplary application followed protocols developed in our lab (22, 23, 41, 18) screening for a) tolerability; b) skin irritation; c) impedance (compliance voltage); and d) material/mechanical properties.

### Device Verification

Electrical performance verification of cell / battery packs under constant current discharge was conducted using a sourcemeter (Keithley 2450 SMU). Electrochemical impedance spectroscopy (EIS) experiments were conducted with a Prinston Applied VersaSTAT 4 Potentiostat/Galvanostat. AC impedance measurements were performed potentiostatically at open circuit voltage and a small signal stimulus of 5 mV within a frequency range of 10 mHz - 10 kHz. The test is conducted with 60 data points distributed across the frequency range on a logarithmic scale. The battery model components were selected based on comparable electrochemistry analysis (26) and fit to EIS data (AMETEK, ZView). Model parameters reflect mass transfer, chemical kinetics, and electrical resistance of current collectors and conductive additives in electrodes, from which device chemistry (particle size, electrode porosity) and structure (thickness of electrode, compaction) can be refined. This includes the upper limit on current density limited by mass transfer of charge carriers. SEM images recorded using SUPRA 55, with an acceleration voltage of 16 kV and backscattered electron detector.

### Load Characterization and Validation

#### Participants

The study was conducted in accordance to protocols and procedures approved by the Institutional Review Board of the City College of New York. All volunteer participants provided written informed consent to participate in the study. The study included 17 subjects (15 male) between the ages of 19 and 38 (M = 25, SD = ±6.4). All subjects were recruited through local advertisement and financially compensated.

#### Screening and exclusion criteria

Participants were excluded if they presented with any skin disorder at or near stimulation locations that compromised skin integrity, such as eczema, rashes, blisters, open wounds, burns including sunburns, cuts, or other skin defects, as the goal of this study was not to determine if skin impairments influence the tolerability or to access electrical stimulation to enhance wound healing.

Stimulation were applied to the ventral or dorsal side of the subject’s left or right forearm. No more than one dose was applied to a region per day (e.g. ventral surface of right arm). All devices were at room temperature (22 °c) immediately prior to testing.

#### Load characterization

For load characterization tests, devices with the same design as Wearable Disposable Electrotherapy but with only interface components and without deposition of active materials were made. Prior to the test, subjects’ forearms were cleansed with soap and water and then dried. After placement of ion conductive hydrogels the monitoring tabs were connected to a sourcemeter (Keithley 2450 SMU). According to the test, the sourcemeter output was set to either constant voltage with a peak current limit, or to constant current with a voltage compliance limit. Output was enabled before placement of the test device on skin.

Temperature was recorded using three thermocouple probes placed in the empty battery pouch, one over the anode stimulation electrode, one over the cathode stimulation electrode and one in the middle of the device over fastening hydrogel. Photographs were taken immediately before and after stimulation under consistent lighting conditions and skin temperature was recorded using a thermal camera (FLIR One Pro). At the beginning and during stimulation, subjects reported subjective pain on a VAS scale every 2 min. Approximately 24 hours after stimulation, subjects’ skin was evaluated for any enduring skin irritation.

#### Validation

The procedure was similar to load tests using the interface test device; instead of connecting the device to the sourcemeter, the Wearable Disposable Electrotherapy Device with active batteries was used. To record the voltage and current of the device, a custom high impedance analog interface was used (0.4 attention factor), a 100 Ω series resistor (for current), and acquisition system (DATAQ DI-1100).

### FEM Device Stimulation

For the exemplary device, we developed a computer-aided design (CAD) model of the Wearable Disposable Electrotherapy prototype and underlying superficial tissue. The biophysical and thermo-electrical properties of biological tissues were based on previous studies and heat-transfer biophysics followed standard assumptions and methods (42, 43). An approximate temperature distribution throughout a perfused tissue can be found by solving the bio-heat transfer (Pennes) equation. For the thermal boundary conditions, all external boundaries were insulated except the top surface which was assigned heat flux. For the tDCS device we used our previously detailed (44) and verified (45) MRI-derived (1 mm^3^ T1/T2; 36 year old male) model. For the iontophoresis device model and the wound healing device model a forearm/hand model was developed from a high-resolution MRI (1.25 mm^3^ T1/T2/Petra; 33 year old male).

CAD structures (devices) were modeled in SolidWorks 2022 (Dassault Systemes Corp., MA, USA) and Simpleware ScanIP U-2022.12-SP2 (Synopsys, WA, USA) imported and numerically solved in COMSOL Multiphysics 5.6 (COMSOL Multiphysics, Boston, MA) under conventional parameters and quasi-static physics (46). The resulting finite element model comprised >32,690,000 tetrahedral elements (>11,120,000 degrees of freedom, with 30 s time-steps) for the exemplary device, >1,880,000 tetrahedral elements (>2,590,000 degrees of freedom) for tDCS model, >50,000,000 tetrahedral elements (>72,000,000 degrees of freedom) for the iontophoresis model, and >35,000,000 tetrahedral elements (>54,000,000 degrees of freedom) for the wound healing model.

## Supplementary data

**Supplementary Fig. 1.**
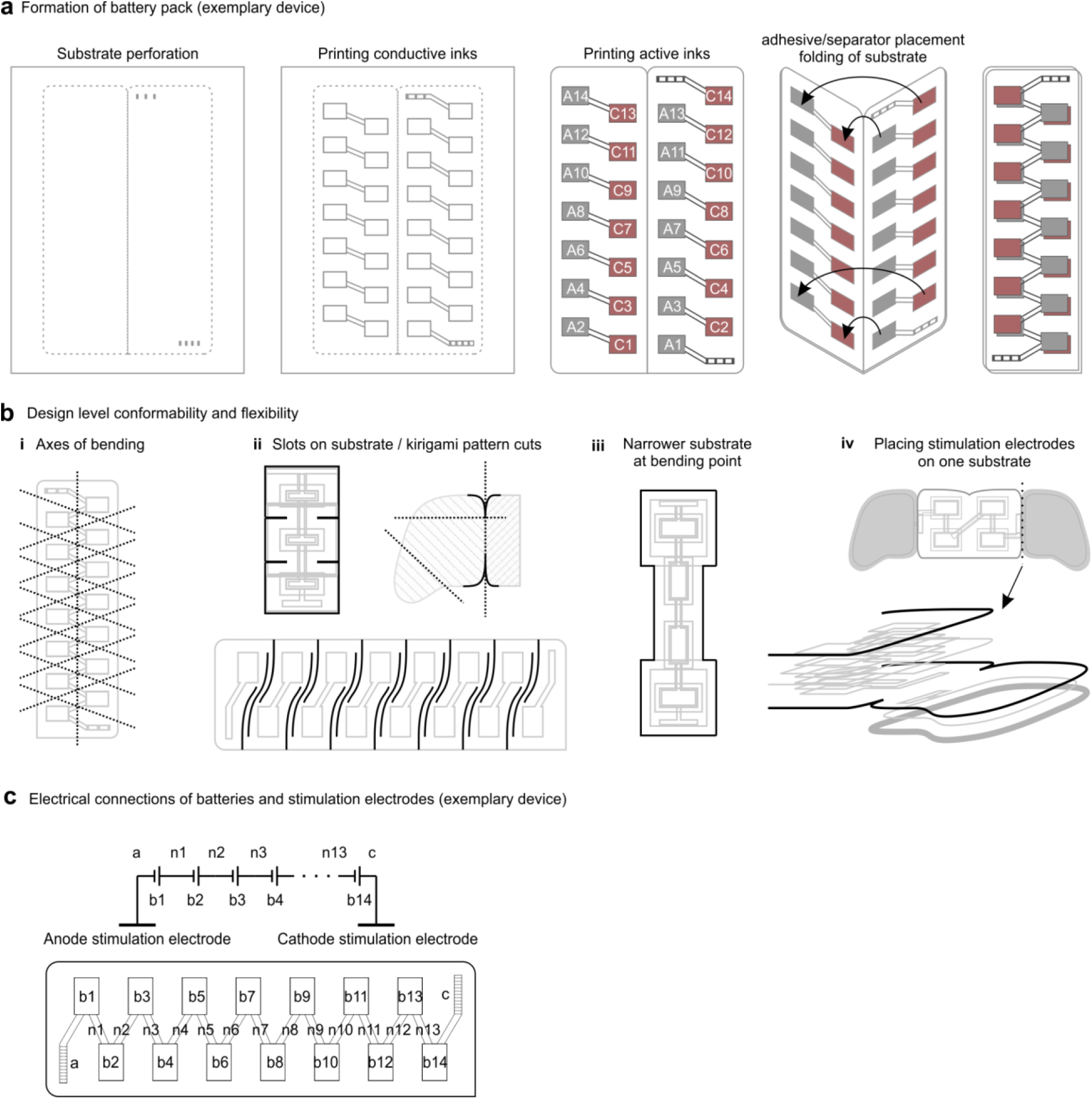
Formation of 3D structure and battery pack functionality of Wearable Disposable Electrotherapy device. **a)** All components, including interconnect and active materials (forming cell pairs, numbered), are printed on a single substrate. In the final manufacturing steps separators and adhesive are added and the device folded, whereby battery cells are formed and connected. **b)** Per application specific requirements, geometry designs further support conformability along low flexural rigidity axes (i), using pattern cuts (ii), and variations in device width (iii) or thickness (iv). **c)** The battery pack output at the terminals (a: stimulation anode, c: stimulation cathode) reflects the series output of battery cells (b1 to b14) through interconnects (n1 to n13).

**Supplementary Fig. 2.**
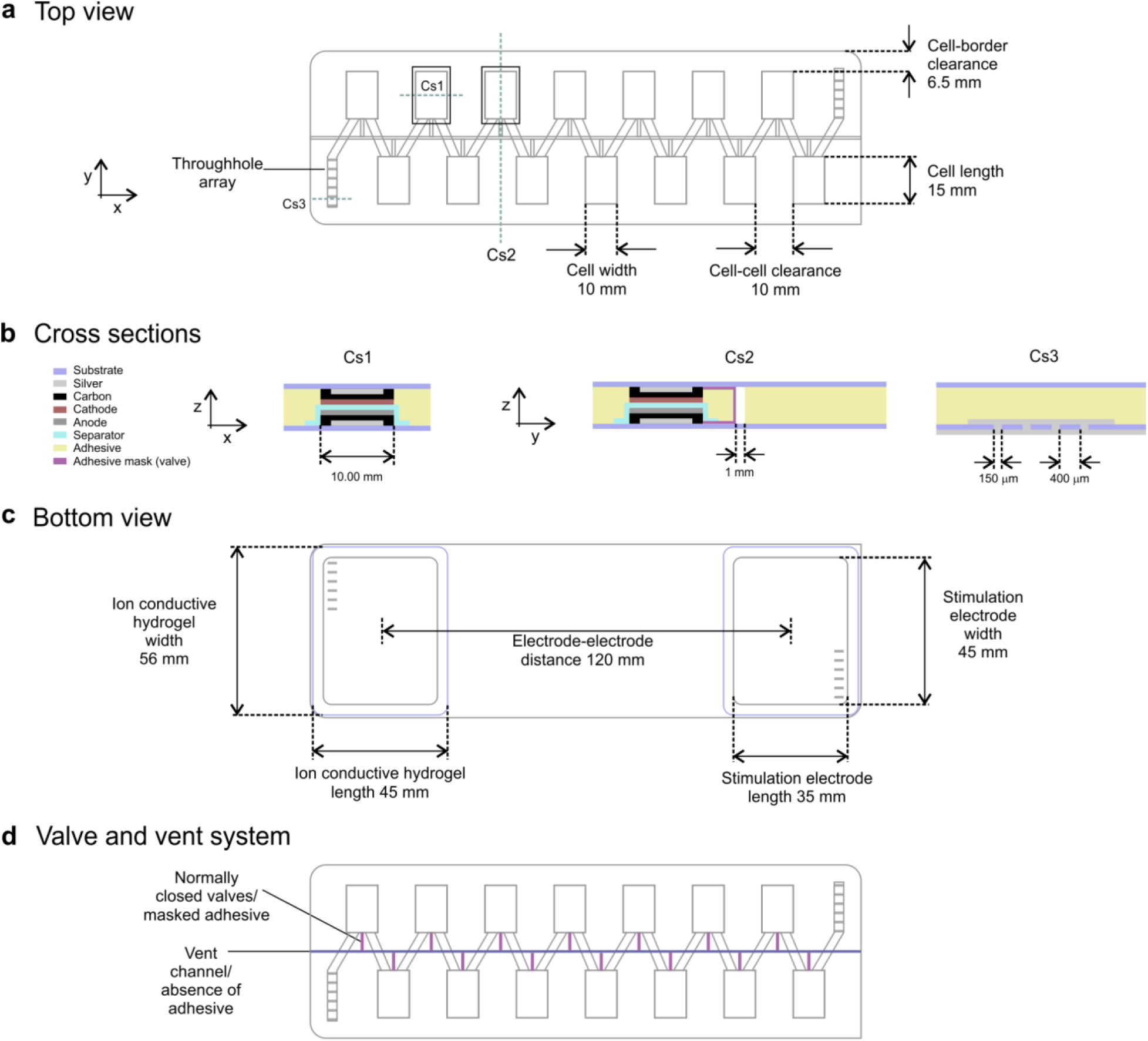
Geometry of exemplary Wearable Disposable Electrotherapy device (battery pack structure and interconnects, interfaces, venting system). **a)** Device top view showing electrochemical architecture of series cells leading to battery pack terminals (with interconnects to bottom substrate), and integrated venting system. **b)** Device cross sections at battery cell (Cs1), at throughholes (Cs3) and at middle of device (Cs2) showing materials and vent channel (false color legend). **c)** Device bottom view showing interface components with throughhole interconnects to top substrate. **d)** Device cross section highlighting venting system.

**Supplementary Fig. 3.**
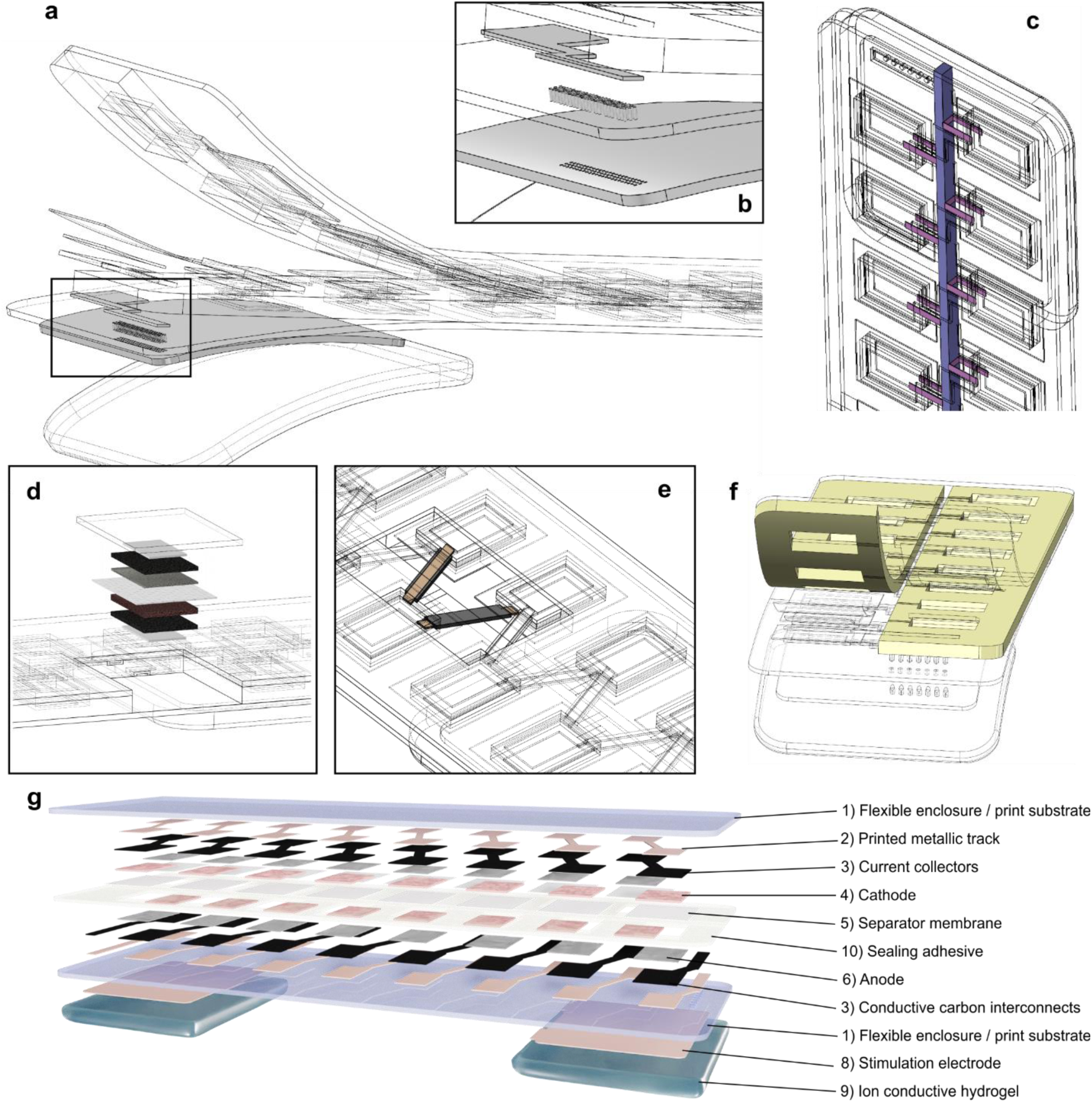
3D Geometry of exemplary Wearable Disposable Electrotherapy device. **a)** Exploded view of device with **b)** Inset showing array of holes on the substrate filled with conductive material, connecting stimulation electrode on back of the substrate to the interconnect on the top of the substrate. **c)** Areal geometry showing the venting system. The vent channel (negative space) is shown in blue and normally closed valves are shown in purple. **d)** Exploded view of layers of one battery cell (top to bottom: substrate enclosure, silver conductor, carbon current collector, anode, separator, cathode, carbon current collector, silver conductor. **e)** View of two interconnects (one on bottom and one on top substrates) connecting one battery cell to two adjacent cells. **f)** Two strips of double-sided tape with cutouts for battery cells, and narrow strips of adhesive masking. The 1 mm gap between adhesives forms the vent channel. **g)** Exploded device view with indexed design elements.

**Supplementary Fig. 4.**
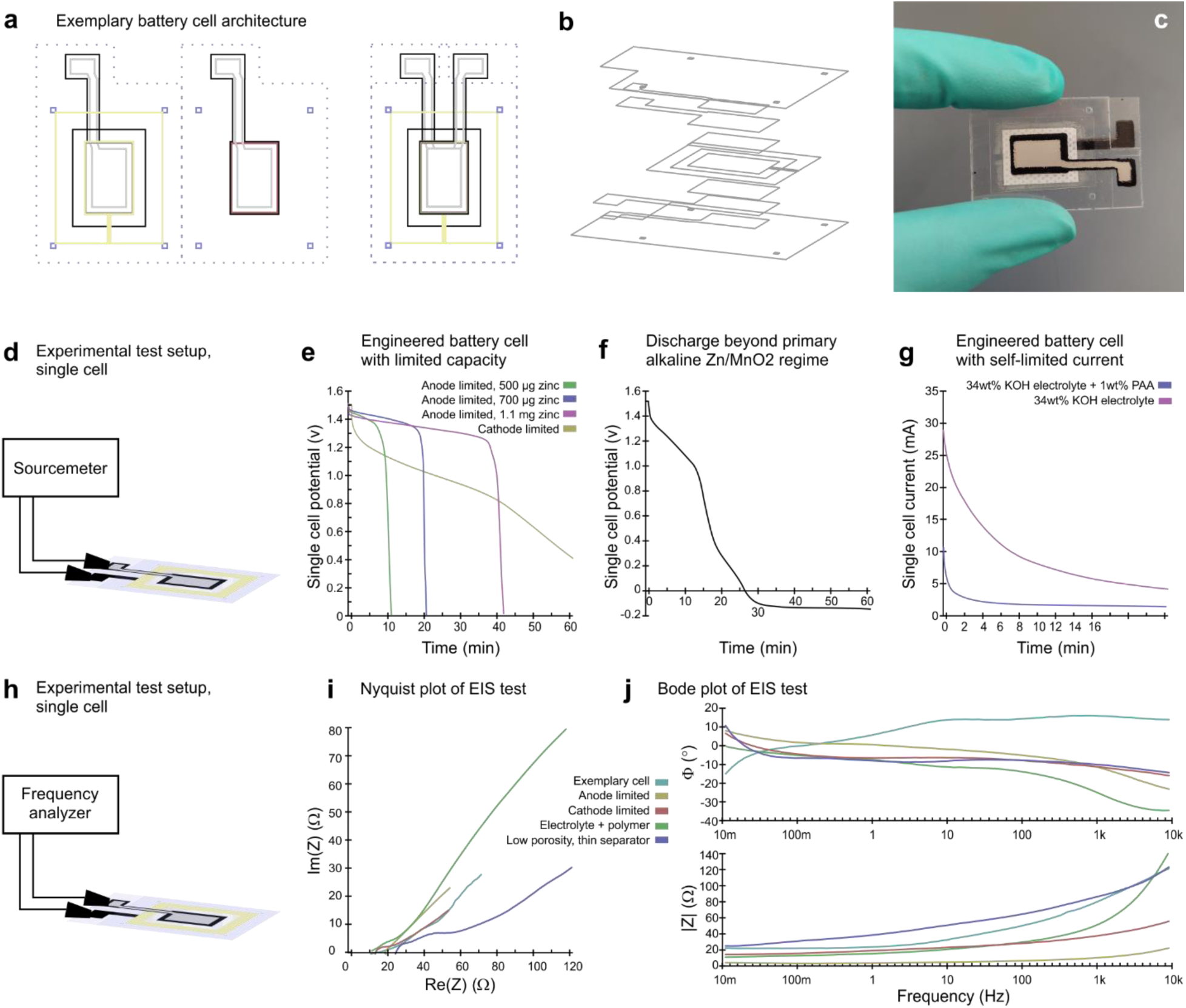
Single battery cell designs (time/current self-limited) and verification. **a)** Design of single packaged cell with interconnects for battery discharge and electrochemical impedance tests. **b)** Exploded view of layers of single cell test device. **c)** Photograph of a single cell test device. **d)** Experimental test setup for a single cell test device discharge test. **e)** Discharge curves for single cells under 700 µA constant current. Note the flat discharge curve followed by sudden drop in anode limited battery vs discharge curve of cathode limited battery. In anode limited batteries, total capacity of the battery is controlled by the mass of zinc. **f)** Discharge curve of a printed battery beyond 0.9 v to predict interaction with load. **g)** Discharge curves for single cells connected to 33 Ω constant load. High current (peak and average) during discharge resulted from a typical cell. Flat discharge and low current rate for a cell with similar mass of active materials with additional current limiting mechanism. Note maximum discharge current can be self-limited independent from load value. **h)** Experimental test setup for a single cell test device electrochemical impedance spectroscopy (EIS). **i)** Nyquist and **j)** Bode plot of electrochemical impedance spectroscopy performed potentiostatically at open circuit voltage with a small signal stimulus of 5 mV across 10 mHz - 10 kHz frequency range.

**Supplementary Figure 5.**
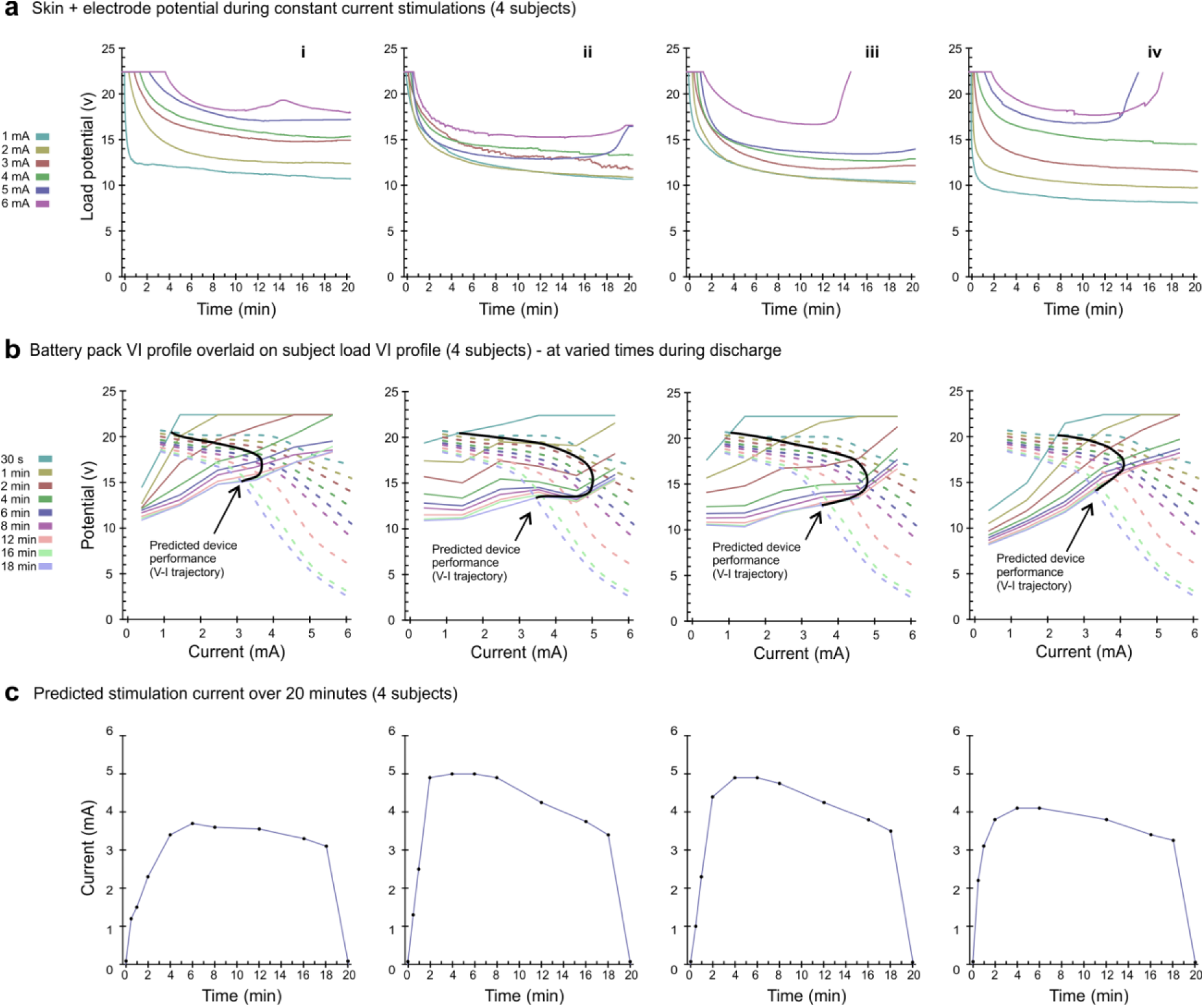
Subject-wise simulation of exemplary Wearable Disposable Electrotherapy device performance. **a)** Interface-physiological load test results for 4 subjects under constant current stimulation at amplitudes of 1-6 mA and voltage limit of 22.4 v during 20 min. **b)** The associated isotemporal V-I profile of subject-specific interface-physiological load results (colored solid lines) are overlaid with battery pack V-I performance (colored dashed lines). Battery pack data is from separately-collected galvanostatic discharge tests (1-6 mA, 20 min). The overlaid interface-physiological load and battery pack V-I profiles are represented for varied time points during discharge (colors). At each timepoint, the intersection of these V-I plots reflects source-load coupling (see theory). The connection of these intersections (solid black line) is then the simulated device discharge trajectory. **c)** This predicted discharge trajectory is shown as current over time performance. On a subject-wise basis, this illustrates the novel design / simulation approach developed for Wearable Disposable Electrotherapy devices, and is then validated.

**Supplementary Figure 6.**
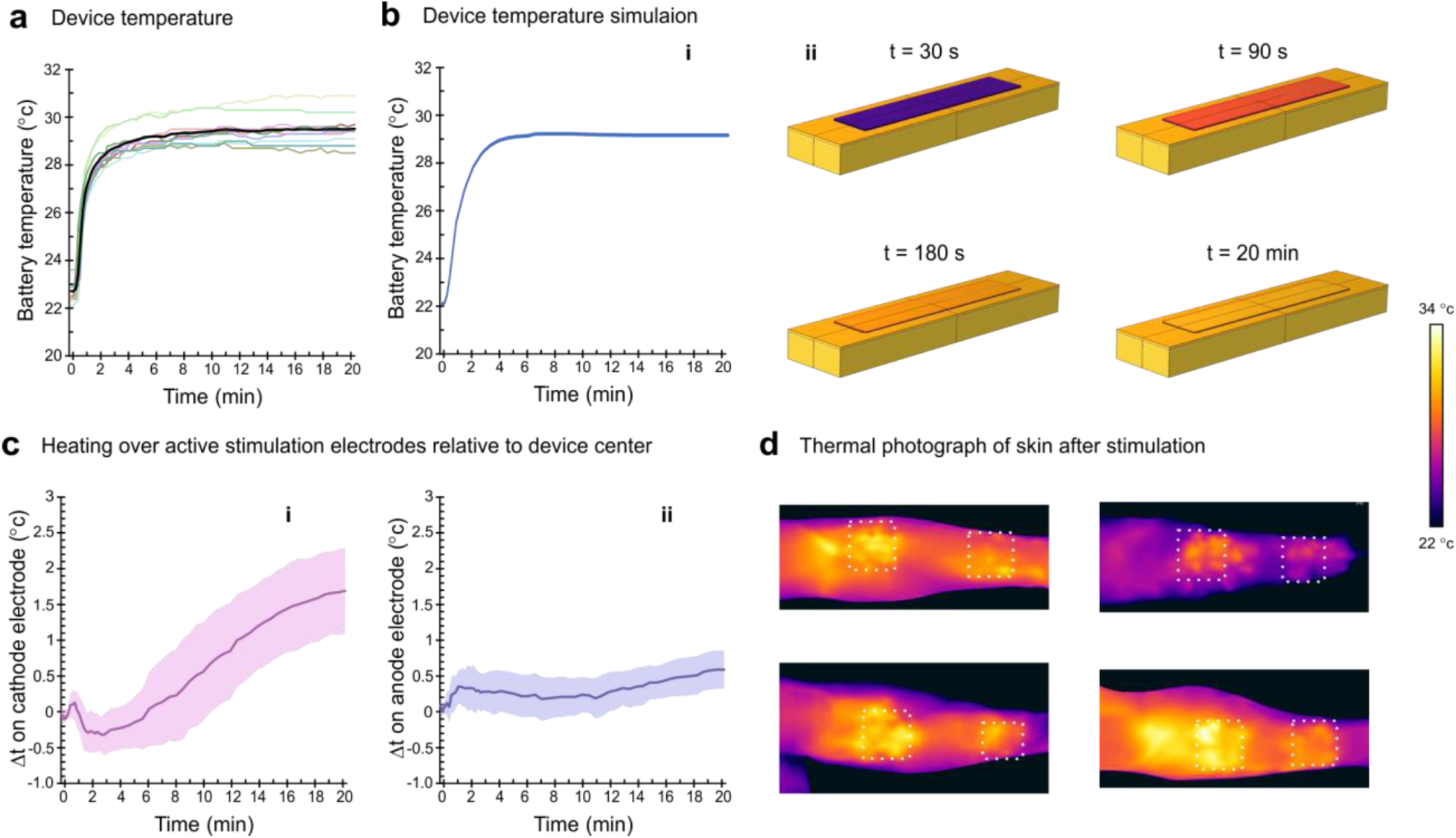
Temperature Transition of exemplary Wearable Disposable Electrotherapy device. Device temperature quickly warms to skin temperature by placement on subjects’ skin (t=0). **a)** Battery pack temperature measured using sensors in place of battery active material in interface test devices for 10 subjects (colored lines) and average (black line). **b)** FEM bioheat simulation of battery pack temperature. **c)** Relative heating of battery material over stimulation cathode electrode (i) and over stimulation anode electrode (ii), normalized to temperature in the center of device (solid lines: average; shaded areas: variance). **d)** IR thermography of skin after stimulation showing relative heating of skin under cathode (left) and anode (right) stimulation electrodes.

**Supplementary Figure 7.**
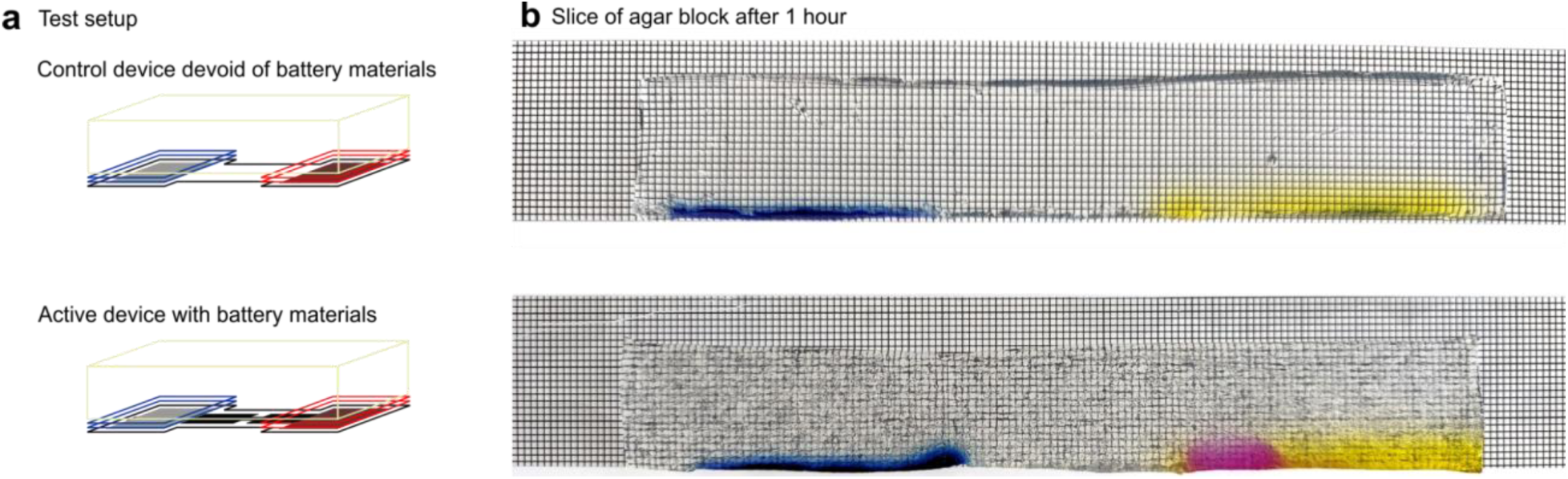
Performance of iontophoresis application Wearable Disposable Electrotherapy device on a skin phantom. (a) Experimental set-up to evaluate enhanced ionic delivery. (b) Top row shows passive diffusion of dyes into phantom placed on the electrodes of a control device (devoid of battery materials). Bottom row shows enhanced dye movement due to iontophoresis, demonstrating the device’s efficacy in facilitating active transport of charged molecules. 0.8 ml of 0.14% Phenol Red (negatively charged, molecular weight 354.4 g/mol) was applied to the nonwoven felt placed on the negative stimulation electrode, and 0.8 ml of 0.2% Methylene Blue (positively charged, molecular weight 319.9 g/mol) was applied to positive stimulation electrode. The phantom blocks were prepared using 250 g of deionized water, 2.5 g of agar powder, and 250 µg of NaCl. The blocks were placed on both the control and active devices for two hours, then sliced and placed on a grid with 1 mm spacing between lines to visualize dye diffusion.

**Supplementary Table 1.**
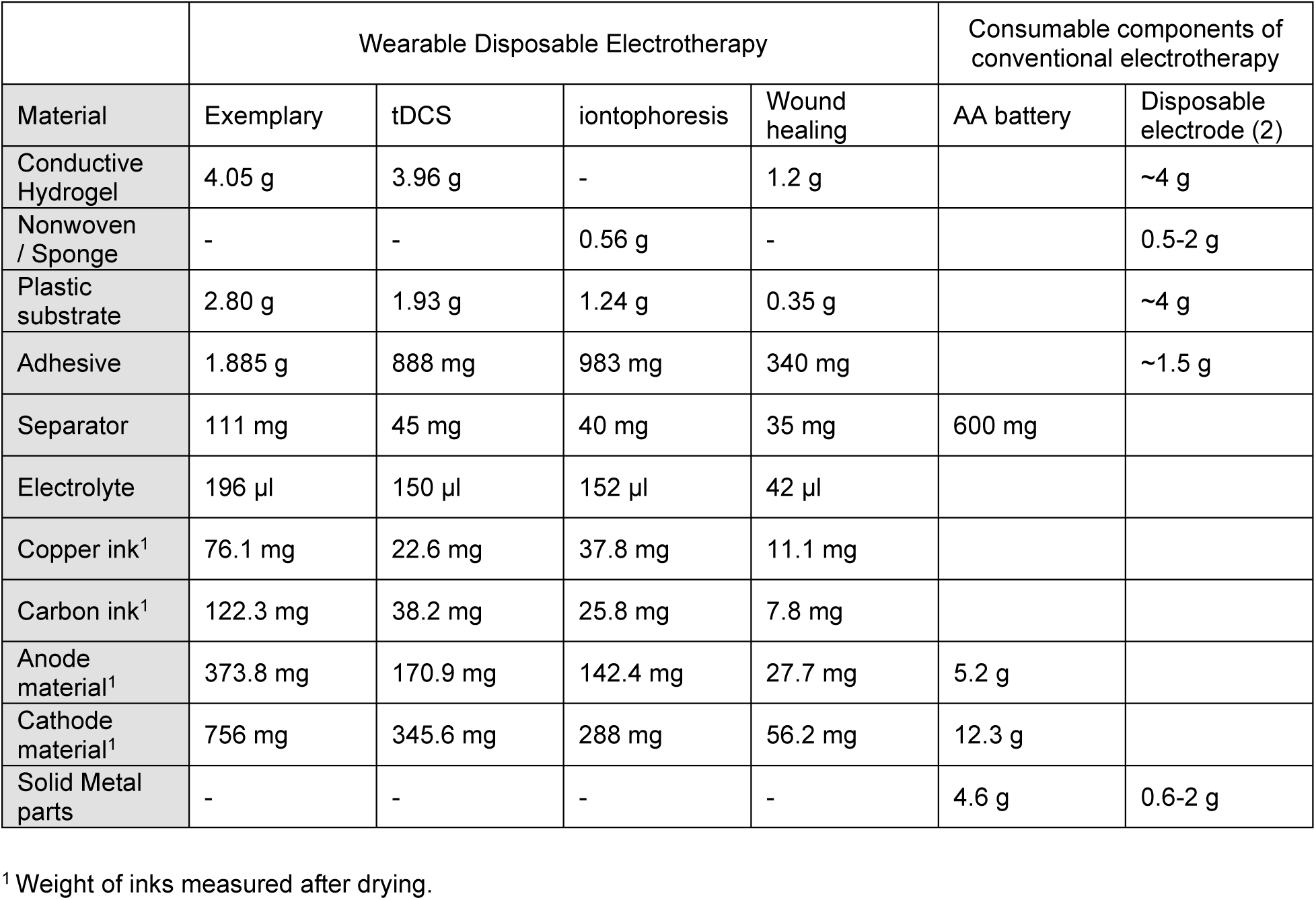
Material use of Wearable Disposable Electrotherapy vs. conventional electrotherapy consumables.

|  | Wearable Disposable Electrotherapy |  |  |  | Consumable components of conventional electrotherapy |  |
| --- | --- | --- | --- | --- | --- | --- |
| Material | Exemplary | tDCS | iontophoresis | Wound healing | AA battery | Disposable electrode (2) |
| Conductive Hydrogel | 4.05 g | 3.96 g | - | 1.2 g |  | ~4 g |
| Nonwoven / Sponge | - | - | 0.56 g | - |  | 0.5-2 g |
| Plastic substrate | 2.80 g | 1.93 g | 1.24 g | 0.35 g |  | ~4 g |
| Adhesive | 1.885 g | 888 mg | 983 mg | 340 mg |  | ~1.5 g |
| Separator | 111 mg | 45 mg | 40 mg | 35 mg | 600 mg |  |
| Electrolyte | 196 µl | 150 µl | 152 µl | 42 µl |  |  |
| Copper ink<sup>1</sup> | 76.1 mg | 22.6 mg | 37.8 mg | 11.1 mg |  |  |
| Carbon ink<sup>1</sup> | 122.3 mg | 38.2 mg | 25.8 mg | 7.8 mg |  |  |
| Anode material<sup>1</sup> | 373.8 mg | 170.9 mg | 142.4 mg | 27.7 mg | 5.2 g |  |
| Cathode material<sup>1</sup> | 756 mg | 345.6 mg | 288 mg | 56.2 mg | 12.3 g |  |
| Solid Metal parts | - | - | - | - | 4.6 g | 0.6-2 g |

<sup>1</sup> Weight of inks measured after drying.

## Supplementary Analysis 1

### 1.0 Isotemporal Trajectory Theoretical Framework Overview

In this section, we characterize the Wearable Disposable Electrotherapy device performance (considered a coupled system of battery pack with interfaces-physiological load) using an approximation of uncoupled parts (Sections 3.0 and 3.1). We base these analyses on the measured behavior of each part (battery and interfaces-physiological load) independently (Section 2.0). These results explain the unique current regulation theory of Wearable Disposable Electrotherapy, as well as the specific simulation step used (Section 4.0) as part of the device design process (Figure 2).

### 2.0 Battery pack and interfaces-physiological load as uncoupled dynamical systems

In our device design approach, we identify and characterize two subsystems: battery pack (B) and interfaces-physiological load (L). First, we consider these two subsystems independently (i.e. without interactions). The voltages *V*_*B*_ and *V*_*L*_ of B and L, respectively, are response functions of time *t*:

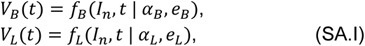

where *I*_*n*_ is a constant current stimulus; *α*_*B*_ and *α*_*L*_ are internal parameters of each system; while *e*_*B*_ and *e*_*L*_ are environmental factors affecting each subsystem. These parameters also have a temporal dependency.

Experimentally, by independently connecting the battery packs or interfaces-physiological load to a source meter, with varied applied constant current levels (e.g. *I*_*n*_ = 1 mA, 2 mA…), and measuring the associated output voltage (*V*_*B*_, *V*_*L*_), the response functions of each system are determined (i.e. *f*_*B*_ and *f*_*L*_).

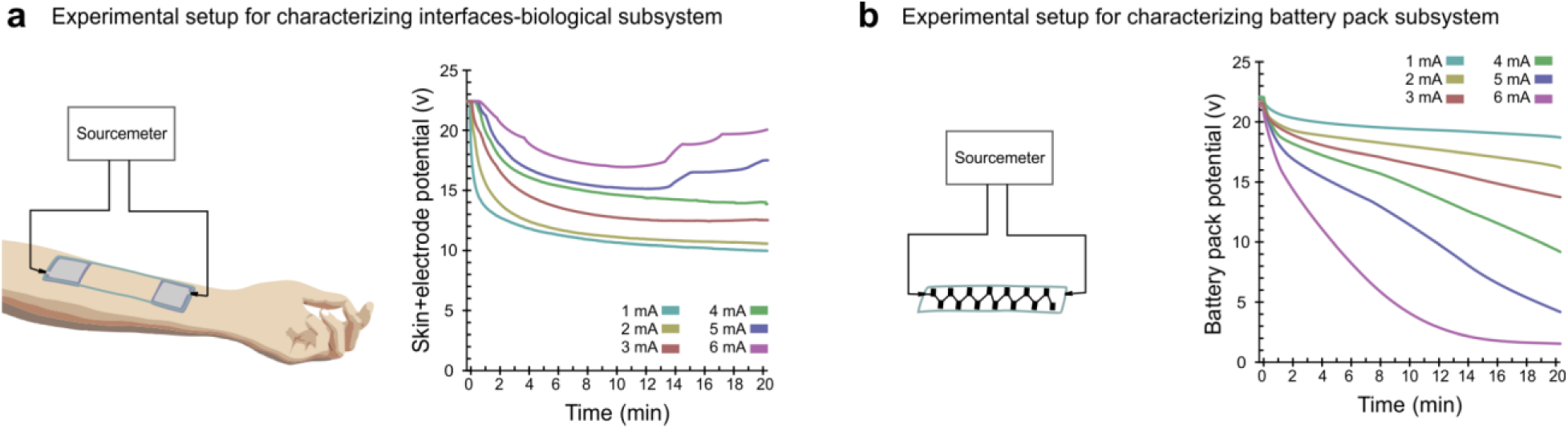

### 3.0 Battery pack and interfaces-physiological load as a coupled dynamical system

In device design two subsystems: battery pack (B) and interfaces-physiological load (L) are in fact physically coupled. Then, system B-L is composed by the subsystems B and L such that both parts exchange the same current *I*(*t*) and have equal voltage *V*(*t*). The voltage of each part of the system evolved with the equations:

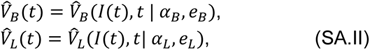

and we know that

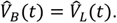

We would like to find solutions to this condition.

### 3.1 Approximation of 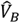 and 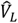 from uncoupled subsystems

Given only independently characterized subsystems (the battery pack and interfaces-physiological load - Section 2.0) our goal is to simulate the performance of the coupled system (ie. the performance of the device when applied to the body).

First, we expand (SA.II) to the first-order Taylor series:

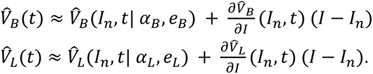

where *I*_*n*_ is a fixed value of *I*(*t*). We denote *t*_0_ any time value such as *I*(*t*_0_) = *I*_*n*_. The condition 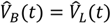 is verified if and only if

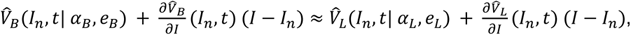

when |*I* − *I*_*n*_| is small.

In the coupled system, the subsystems exchange I(t). At time t_0_, the value of I(t_0_) is I_n_. At this particular instant, one can suppose systems B and L receive I_0_ independently (i.e. as if they were two uncoupled systems - Section 2.0).

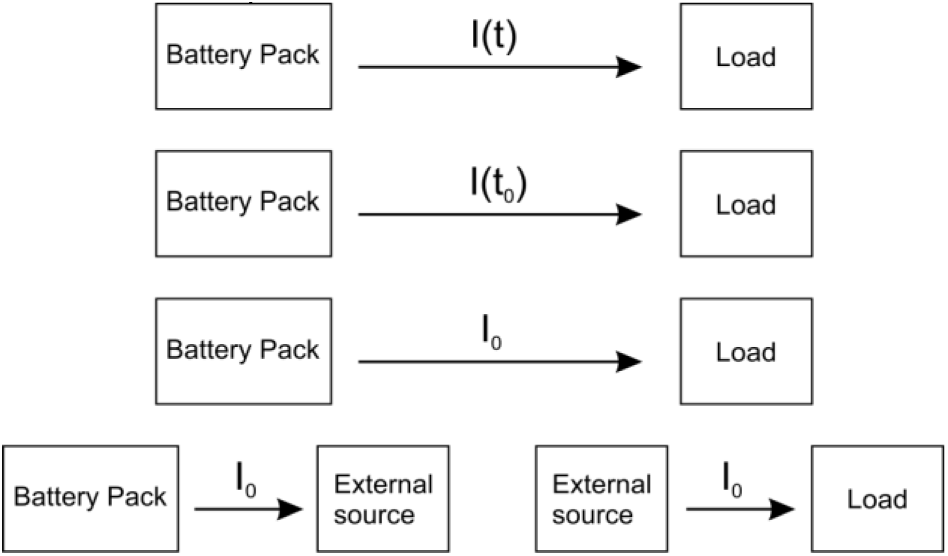

Then,

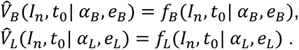

In summary,

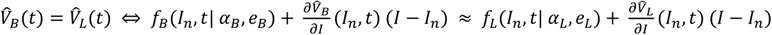

Moreover, since |*I* − *I*_*n*_|) is small, we can reduce it to

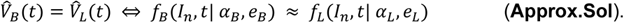

Therefore the approximate solution to the coupled system is described by the response functions under fixed constant current of the uncoupled system. Indeed, these response functions are known (from 2.0)! We assume the environmental dependencies (*e*_*B*_ and *e*_*L*_) are comparable under experiment-isolated system testing and couple system discharge.

(**Approx Sol**) suggest we can solve the coupled system (B-L) performance when a device is applied to the body - by calculating the intersection of the response functions *f*_*B*_ and *f*_*L*_ - which are measured as independent subsystems (B and L):

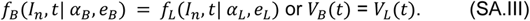

In the next section, we show the numerical implementation of this solution.

### 4.0 Numerical solution of simulated coupled subsystems

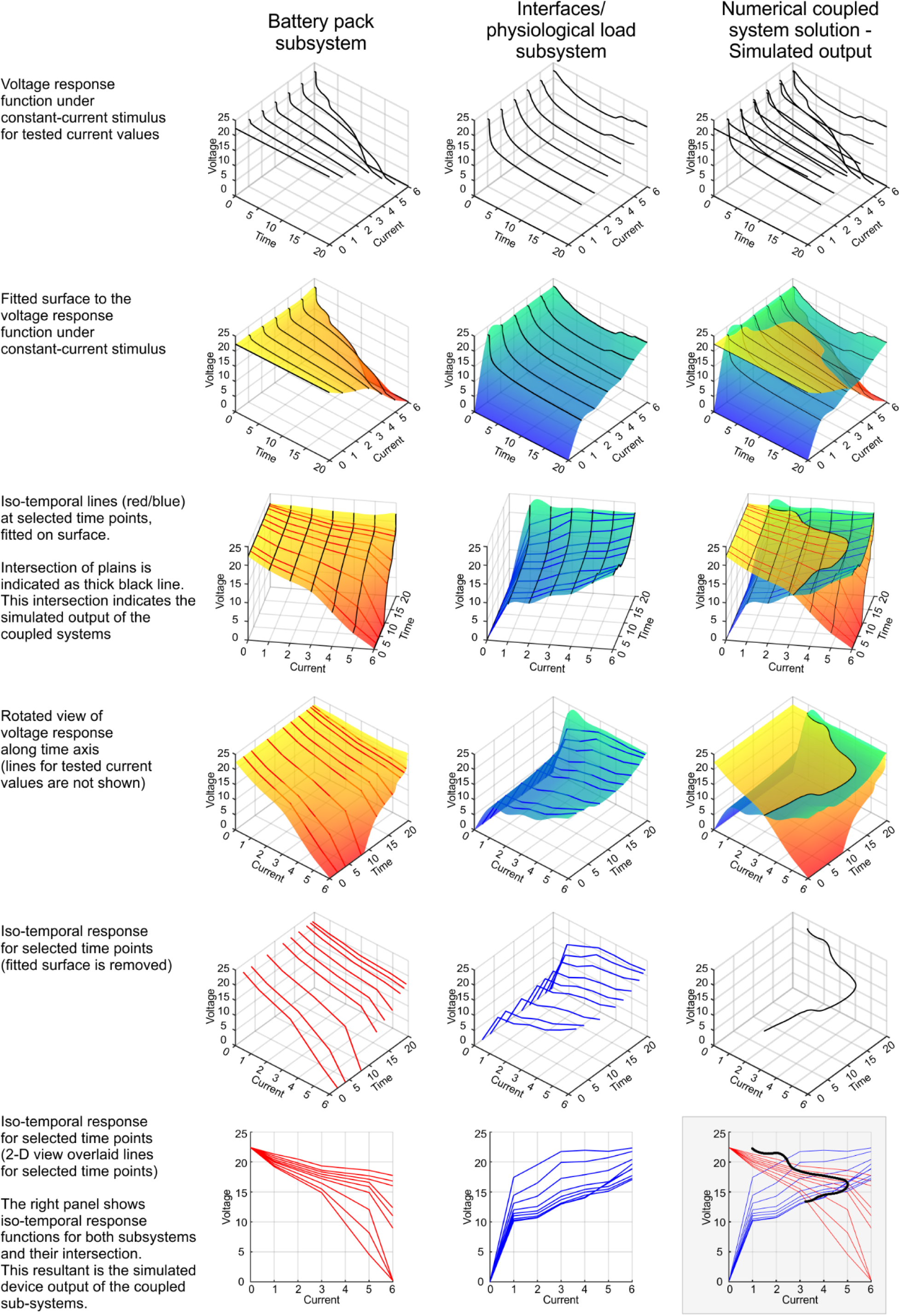

## Notes

### Competing Interest Statement

The City University of New York holds patents on brain stimulation or battery technology with MB, AC, MFR as inventor. MB has equity in Soterix Medical Inc. AC has equity in Urban Electric Power. MB consults, received grants, assigned inventions, and/or served on the SAB of SafeToddles, Boston Scientific, GlaxoSmithKline, Biovisics, Mecta, Lumenis, Halo Neuroscience, Google-X, i-Lumen, Humm, Allergan (Abbvie), Apple, Ybrain, Ceragem, Remz. MB is supported by grants from Harold Shames and the National Institutes of Health: NIH-NIDA UG3DA048502, NIH-NIGMS T34 GM137858, NIH-NINDS R01 NS112996, NIH-NINDS R01 NS101362, and NIH-G-RISE T32GM136499.

### Summary of Updates

Illustrated design and optimization pipeline. Show 3 application examples for Wearable Disposable Electrotherapy platform. Additional human trials for new applications. Enhanced theoretical background for complex interaction between device and biological load.

## References

1. Vance, C. G., Dailey, D. L., Rakel, B. A. & Sluka, K. A. Using TENS for pain control: the state of the evidence. Pain management 4, 197–209 (2014).

2. Coppola, G. et al. Neuromodulation for chronic daily headache. Current Pain and Headache Reports 26, 267–278 (2022).

3. Fregni, F. et al. Evidence-based guidelines and secondary meta-analysis for the use of transcranial direct current stimulation in neurological and psychiatric disorders. International Journal of Neuropsychopharmacology 24, 256–313 (2021).

4. Antal, A. et al. Low intensity transcranial electric stimulation: safety, ethical, legal regulatory and application guidelines. Clinical neurophysiology 128, 1774–1809 (2017).

5. Thakral, G. et al. Electrical stimulation to accelerate wound healing. Diabetic foot & ankle 4, 22081 (2013).

6. Jiang, Y. et al. Wireless, closed-loop, smart bandage with integrated sensors and stimulators for advanced wound care and accelerated healing. Nature biotechnology 41, 652– 662 (2023).

7. Bikson, M. et al. Limited output transcranial electrical stimulation (LOTES-2017): Engineering principles, regulatory statutes, and industry standards for wellness, over-the-counter, or prescription devices with low risk. Brain stimulation 11, 134–157 (2018).

8. Karpiński, T. M. Selected medicines used in iontophoresis. Pharmaceutics 10, 204 (2018).

9. Wang, W. et al. Neuromorphic sensorimotor loop embodied by monolithically integrated, low-voltage, soft e-skin. Science 380, 735–742 (2023).

10. Jiang, M., Lu, Y., Zhu, Z. & Jia, W. Advances in Smart Sensing and Medical Electronics by Self-Powered Sensors Based on Triboelectric Nanogenerators. Micromachines 2021 12, 698. (2021).

11. Guleyupoglu, B., Schestatsky, P., Edwards, D., Fregni, F. & Bikson, M. Classification of methods in transcranial electrical stimulation (tES) and evolving strategy from historical approaches to contemporary innovations. Journal of neuroscience methods 219, 297–311 (2013).

12. Dhote, V., Bhatnagar, P., Mishra, P. K., Mahajan, S. C. & Mishra, D. K. Iontophoresis: a potential emergence of a transdermal drug delivery system. Scientia pharmaceutica 80, 1–28 (2012).

13. Peebles, I. S., Phillips, T. O. & Hamilton, R. H. Toward more diverse, inclusive, and equitable neuromodulation. Brain Stimulation 16, 737–741 (2023).

14. Ye, C. et al. A wearable aptamer nanobiosensor for non-invasive female hormone monitoring. Nature Nanotechnology 19, 330–337 (2024).

15. Xu, G. et al. Battery-free and wireless smart wound dressing for wound infection monitoring and electrically controlled on-demand drug delivery. Advanced Functional Materials 31, 2100852 (2021).

16. Peterchev, A. V. et al. Fundamentals of transcranial electric and magnetic stimulation dose: definition, selection, and reporting practices. Brain stimulation 5, 435–453 (2012).

17. Merrill, D. R., Bikson, M. & Jefferys, J. G. Electrical stimulation of excitable tissue: design of efficacious and safe protocols. Journal of neuroscience methods 141, 171–198 (2005).

18. Minhas, P. et al. Electrodes for high-definition transcutaneous DC stimulation for applications in drug delivery and electrotherapy, including tDCS. Journal of neuroscience methods 190, 188–197 (2010).

19. Gaikwad, A. M., Steingart, D. A., Ng, T. a N., Schwartz, D. E. & Whiting, G. L. A flexible high potential printed battery for powering printed electronics. Applied Physics Letters 102, (2013).

20. Newman, J. S. & Tobias, C. W. Theoretical analysis of current distribution in porous electrodes. Journal of The Electrochemical Society 109, 1183 (1962).

21. Lanzi, O. & Landau, U. Effect of pore structure on current and potential distributions in a porous electrode. Journal of the Electrochemical Society 137, 585 (1990).

22. Paneri, B. et al. Tolerability of repeated application of transcranial electrical stimulation with limited outputs to healthy subjects. Brain Stimulation 9, 740–754 (2016).

23. Khadka, N. et al. Dry tDCS: Tolerability of a novel multilayer hydrogel composite non-adhesive electrode for transcranial direct current stimulation. Brain Stimulation 11, 1044–1053 (2018).

24. Bora, D. J. & Dasgupta, R. Estimation of skin impedance models with experimental data and a proposed model for human skin impedance. IET Systems Biology 14, 230–240 (2020).

25. Unal, G. et al. Quasi-static pipeline in electroconvulsive therapy computational modeling. Brain stimulation 16, 607–618 (2023).

26. Lazanas, A. C. & Prodromidis, M. I. Electrochemical impedance spectroscopy─ a tutorial. ACS Measurement Science Au. 3, 162–193 (2023).

27. Khadka, N. et al. Minimal heating at the skin surface during transcranial direct current stimulation. Neuromodulation: Technology at the Neural Interface 21, 334–339 (2018).

28. Xu, X., Zhang, H., Yan, Y., Wang, J. & Guo, L. Effects of electrical stimulation on skin surface. Acta Mechanica Sinica 1–29 (2021).

29. Ezquerro, F. et al. The influence of skin redness on blinding in transcranial direct current stimulation studies: a crossover trial. Neuromodulation: Technology at the Neural Interface 20, 248–255 (2017).

30. Liu, S. et al. Conformability of flexible sheets on spherical surfaces. Science Advances 9, eadf2709 (2023).

31. Woods, A. J. et al. A technical guide to tDCS, and related non-invasive brain stimulation tools. Clinical neurophysiology 127, 1031–1048 (2016).

32. Seibt, O., Brunoni, A. R., Huang, Y. & Bikson, M. The pursuit of DLPFC: non-neuronavigated methods to target the left dorsolateral pre-frontal cortex with symmetric bicephalic transcranial direct current stimulation (tDCS). Brain stimulation 8, 590–602 (2015).

33. Dixit, N., Bali, V., Baboota, S., Ahuja, A. & Ali, J. Iontophoresis-an approach for controlled drug delivery: a review. Current Drug Delivery 4, 1–10 (2007).

34. Guest, J. F., Singh, H., Rana, K. & Vowden, P. Cost-effectiveness of an electroceutical device in treating non-healing venous leg ulcers: results of an RCT. Journal of Wound Care 27, 230–243 (2018).

35. Hao, Q., Horton, J. & Hamson, A. Electrostimulation Devices for Wounds. Canadian Journal of Health Technologies 3, (2023).

36. Karpiński, T. M. Selected medicines used in iontophoresis. Pharmaceutics 10, 204 (2018).

37. Simpson, K. N., Welch, M. J., Kozel, F. A., Demitrack, M. A. & Nahas, Z. Cost-effectiveness of transcranial magnetic stimulation in the treatment of major depression: a health economics analysis. Advances in Therapy 26, 346–368 (2009).

38. Leiphart, J., Barrett, M. & Shenai, M. B. Economic inequities in the application of neuromodulation devices. Cureus 11, (2019).

39. Kadakia, K. T. et al. Challenges and solutions to advancing health equity with medical devices. Nature Biotechnology 41, 607–609 (2023).

40. Sousa, R. E., Costa, C. M. & Lanceros-Méndez, S. Advances and future challenges in printed batteries. ChemSusChem. 8, 3539–3555 (2015).

41. Guleyupoglu, B., Febles, N., Minhas, P., Hahn, C. & Bikson, M. Reduced discomfort during high-definition transcutaneous stimulation using 6% benzocaine. Frontiers in Neuroengineering 7, 28 (2014).

42. Zannou, A. L., Khadka, N. & Bikson, M. Bioheat model of spinal column heating during high-density spinal cord stimulation. Neuromodulation: Technology at the Neural Interface 26, 1362–1370 (2023).

43. Datta, A., Elwassif, M. & Bikson, M. Bio-heat transfer model of transcranial DC stimulation: comparison of conventional pad versus ring electrode. 2009 Annual International Conference of the IEEE Engineering in Medicine and Biology Society 670–673 (2009).

44. Datta, A., Truong, D., Minhas, P., Parra, L. C. & Bikson, M. Inter-individual variation during transcranial direct current stimulation and normalization of dose using MRI-derived computational models. Frontiers in Psychiatry 3, 91 (2012).

45. Huang, Y. et al. Measurements and models of electric fields in the in vivo human brain during transcranial electric stimulation. Elife. 6, e18834 (2017).

46. Wang, B. et al. Quasistatic approximation in neuromodulation. arXiv preprint arXiv:2402.00486 (2024).

